# Genetic Architecture and Analysis Practices of Circulating Metabolites in the NHLBI Trans-Omics for Precision Medicine (TOPMed) Program

**DOI:** 10.1101/2024.07.23.604849

**Authors:** Nannan Wang, Franklin P. Ockerman, Laura Y. Zhou, Megan L. Grove, Taryn Alkis, John Barnard, Russell P. Bowler, Clary B. Clish, Shinhye Chung, Emily Drzymalla, Anne M. Evans, Nora Franceschini, Robert E. Gerszten, Madeline G. Gillman, Scott R. Hutton, Rachel S. Kelly, Charles Kooperberg, Martin G. Larson, Jessica Lasky-Su, Deborah A. Meyers, Prescott G. Woodruff, Alexander P. Reiner, Stephen S. Rich, Jerome I. Rotter, Edwin K. Silverman, Vasan S. Ramachandran, Scott T. Weiss, Kari E. Wong, Alexis C. Wood, Lang Wu, NHLBI Trans-Omics for Precision Medicine (TOPMed) Consortium, Ronit Yarden, Thomas W. Blackwell, Albert V. Smith, Han Chen, Laura M. Raffield, Bing Yu

## Abstract

Circulating metabolite levels partly reflect the state of human health and diseases and can be impacted by genetic determinants. Hundreds of loci associated with circulating metabolites have been identified; however, most findings focus on predominantly European ancestry or single-study analyses. Leveraging the rich metabolomics resources generated by the NHLBI Trans-Omics for Precision Medicine (TOPMed) Program, we harmonized and accessibly cataloged 1,729 circulating metabolites among 25,058 ancestrally diverse samples. We provided a set of reasonable strategies for outlier and imputation handling to process metabolite data. Following the practical analysis framework, we further performed a genome-wide association analysis on 1,135 selected metabolites using whole genome sequencing data from 16,359 individuals passing the quality control filters, and discovered 1,778 independent loci associated with 667 metabolites. Among 108 novel locus-metabolite pairs, we detected not only novel loci within previously implicated metabolite associated genes but also novel genes (such as *GAB3* and *VSIG4* located in the X chromosome) that have putative roles in metabolic regulation. In the sex-stratified analysis, we revealed 85 independent locus-metabolite pairs with evidence of sexual dimorphism, including well-known metabolic genes such as *FADS2*, *D2HGDH*, *SUGP1*, *UTG2B17*, strongly supporting the importance of exploring sex difference in the human metabolome. Taken together, our study depicted the genetic contribution to circulating metabolite levels, providing additional insight into the understanding of human health.

## Introduction

Metabolites, small molecules produced during metabolism, play a pivotal role in cellular and physiological processes. They are widely used as clinical biomarkers, as their observed levels in biofluids can reflect physiological and pathophysiological processes. Many metabolites are highly heritable^1,2^, indicating a strong genetic influence. Investigating the genetic underpinnings of circulating metabolite levels, which requires rigorous data quality control and integration due to diverse technologies used for measurement, can enhance our understanding of the pathways involved in diseases and health.

The pre-processing and integration of metabolomic data presents significant challenges. Consistent protocols for data integration are essential to maximize the potential of the metabolomic data resource. Investigators face several critical challenges. Outliers, often biologically plausible, can be more difficult to handle in standard linear modeling analysis methods. Common strategies for handling outliers in metabolomic data include extreme value removal^3,4^, inverse normal transformation (INT)^5,6^, and winsorization^7^, each with trade-offs in terms of data distribution and potential loss of information. Meta-analysis^8–12^ is frequently used for the integration of association evidence from multiple studies. This approach allows for study-specific considerations and addresses privacy concerns when individual-level data cannot be directly combined in a pooled analysis. However, meta-analyses can be prone to biases and reduced statistical power, particularly when detecting individual-level interactions or in the analysis of rare variants. Pooling metabolomics data across studies can increase statistical power^13^, but introduce challenges such as platform variability and missing data patterns among studies. Missing data imputation is another critical challenge. The simple imputation to a single value and more sophisticated methods, such as Random Forest^14,15^, and k-Nearest Neighbors (KNN)^16^, are frequently used for imputation.

Genome-wide associations with the metabolome have been reported^6,17–24^. Most previous studies focused on a single specific population, such as European origin^17,18,22,24^, Hispanics^20^, Israeli^21^, Black^23^. Some studies enroll the samples from a single study^17,18,22^. Genetic factors influencing metabolism can vary significantly between different ethnic groups, thus combining data from multiple populations and multiple studies can better capture the full range of genetic variation across human populations. Moreover, research suggests that significant differences may exist between males and females in the incidence or severity of diseases, metabolism, and reaction to medications^25–27^. Although sex dimorphisms on several metabolites have been reported^28–30^, metabolome-wide sex-stratified analyses have not been performed on the genome-wide scale to systematically investigate the genetic architecture of circulating metabolites that differs by sex.

The National Heart Lung and Blood Institute (NHLBI) Trans-Omics for Precision Medicine (TOPMed) program aims to provide information that allows the personalization of disease treatments based on an individual’s unique genes and environment. TOPMed is generating a rich resource of metabolomic data across multiple studies at two TOPMed Centralized Omics Resource Core (CORE) centers: Baylor□UTHealth Metabolomics Center with Metabolon, Inc., and Broad-Beth Israel Deaconess Medical Center; hereafter referred to as the Global Discovery Panel and Broad-BIDMC Panel, respectively. Our study harmonized and cataloged the circulating metabolites measured from eight studies at the two TOPMed CORE centers, with participants from diverse populations. We then explore the impacts of different metabolomic data pre-processing strategies, i.e., outlier handling, missingness imputation, multi-studies analysis strategies, on the association analysis of age and sex with metabolites, providing a practical analysis framework for metabolite data pre-processing. Following the above practice analysis framework, we conducted association analyses using whole genome sequence data (WGS) to investigate the associations with 1,135 selected metabolites. We also systematically assessed sex differences in the genetic regulation of the circulating metabolites. Our findings from genome-wide association and sex-stratified analyses will improve our understanding on the genetic architecture of circulating metabolites and advance our knowledge on their physiological and pathophysiological processes.

## Subjects and Methods

### Metabolite Data Acquisition

Metabolite data for eight studies were generated at the following two TOPMed CORE centers: 1) the Global Discovery Panel was used to process the Genetic Epidemiology of Chronic Obstructive Pulmonary Disease (COPDGene), the Subpopulations and Intermediate Outcomes in Chronic Obstructive Pulmonary Disease Study (SPIROMICS) and Women’s Health Initiative (WHI); and 2) the Broad-BIDMC Panel was used in Childhood Asthma Management Program (CAMP), Genetic Epidemiology of Asthma in Costa Rica (CRA), Framingham Heart Study (FHS), Multi-Ethnic Study of Atherosclerosis (MESA), and WHI. WHI had metabolites measured at both COREs and considered as two studies, i.e., WHI-Baylor-UTHealth and WHI-Broad-BIDMC. All studies used plasma derived from venous blood.

The Global Discovery Panel is comprised of multiple chromatographic methods encompassing an ACQUITY reverse phase ultra-performance liquid chromatography tandem mass spectrometry (RP/UPLC-MS) (Waters; Milford, MA) and a Q Exactive accurate-mass liquid chromatography mass spectrometer (LC-MS) (Thermo Fisher Scientific; Waltham, MA) interfaced with a heated electrospray ionization (HESI-II) source, and Orbitrap mass analyzer operated at 35,000 mass resolution. Raw mass spectrometry data were analyzed using custom developed software (Metabolon, Inc.; Morrisville, NC) and compounds were identified by comparing them to library entries of purified standards or recurrent unknown entities. Metabolon maintains a library based on the retention time index (RI), mass to charge ratio (m/z), and chromatographic data established from >5,400 authenticated standards. Detailed methods were previously described^31–34^.

Metabolomic profiles by the Broad-BIDMC Panel were obtained using liquid chromatography mass spectrometry (LC-MS) as previously described^35,36^. In brief, water-soluble, polar metabolites were assayed with a Nexera X2 U-HPLC (Shimadzu Corp; Marlborough, MA) using an Atlantis hydrophilic interaction LC column (Waters) coupled to a Q Exactive hybrid quadrupole Orbitrap mass spectrometer (Thermo Fisher Scientific). Raw data were visualized using TraceFinder 3.1 (Thermo Fisher Scientific) and Progenesis QI (Nonlinear Dynamics; Newcastle upon Tyne, United Kingdom). A 1290 Infinity LC system (Agilent Technologies) with an XBridge Amide column (Waters) was used in negative ionization or amide mode to measure organic acids, sugars, purines, pyrimidines, and other intermediary metabolites. The LC was coupled to a 6490 Triple Quad MS (Agilent Technologies) in multiple reaction monitoring (MRM) scanning mode, and MassHunter Quantitative Analysis software V10.1 (Agilent Technologies) was used for peak identification.

### Metabolite Harmonization and Catalog

Study specific metabolite data was obtained through dbGaP across eight studies from the two TOPMed CORE centers. Metabolite names and available identifiers, such as Human Metabolome Database (HMDB), KEGG, and chemical IDs (unique to the Global Discovery Panel), were provided in metadata from each data generating core across studies. We used HMDB and metabolite names based on the latest Broad metabolite library, as well as manual check, to harmonize across studies profiled by the Broad Institute. Metabolites profiled by the Broad-BIDMC Panel were measured using multiple Broad methods, and we used a priority of C8-pos, HILIC-pos, C18-neg, and Amide (highest to lowest) to only keep unique metabolites. This priority was based on the metabolomic core recommendation and the metabolite missing rates and CV values across Broad-BIDMC method studies. To harmonize across studies profiled by Global Discovery Panel, we used Metabolon chemical IDs, HMDB, and metabolite names based on the latest Metabolon metabolite library. Metabolites provided by the Global Discovery Panel were based on internal platform priority. To harmonize metabolites across metabolomic cores, we used the HMDB and metabolite names provided by the RefMet workbench and confirmed with representatives at each metabolomic core. Duplicate metabolites were identified as having the same metabolite name or same HMDB, and unique metabolites were retained based on method priority and missing rates. After harmonization, the available RefMet name provided by Metabolomics Workbench was assigned to each metabolite to produce a cross-platform and cross-study metabolite catalog. We note that our analyses here prioritize named metabolites only, unknown metabolites are also available in TOPMed datasets but are not categorized and assessed in the current work due to incomplete chemical structural information.

### Pre-processing metabolites

#### Treatment of Metabolite Outliers

Three methods were considered in our study, including removing observations more than 6 standard deviations away from the mean metabolite value (6SD), within-cohort rank-based INT, and winsorization, where outliers are replaced by the percentile value. In winsorization, for observations above the 95th percentile or below the 5th percentile, we replaced them with the 95th percentile and 5th percentile, respectively. All three methods were initially applied within studies for each metabolite, removing missing observations. For comparability, we then apply INT to the pooled data across studies. To reduce the computational burden, and in concordance with prior work using TOPMed study generated data, we compare the outlier methods using pooled data.

#### Pooled vs. Meta-Analysis

To assess the best method in TOPMed for such cross-study analysis, we compared a meta-analysis versus pooling across the studies. Due to the different platforms and studies, not all metabolites will be measured in every study. To compare meta-analysis with pooled analysis, we only considered metabolites that have measurements in 2 or more studies. We used complete-case with no missing imputation and compared the parameter estimate as well as the *P* value in the meta-analysis versus pooled analysis using 6SD, INT, or winsorization. To compare these strategies, we examined age and sex associations with metabolites. We removed related individuals. FHS was the only study with significant numbers of related individuals and using the pedigree file we removed individuals from the same family.

In a pooled analysis, for the *i*-th metabolite, we fit the age association model

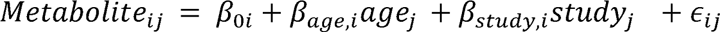

Where *j* denotes the subject with measurement of the *i*-th metabolite and *study_j_* is a categorical variable.

In a meta-analysis, we used the inverse variance-weighted (IVW) method to perform fixed effects meta-analysis. IVW calculates the weighted mean of the effect size estimates using the inverse variance of the individual studies as weights to summarize effect sizes from the independent studies. This model assumes that studies share a single true effect size. Thus, for *i*-th metabolite, we fit the age association model

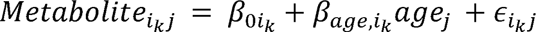

To integrate multiple effect size estimates from multiple studies, we estimated 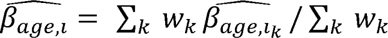, where *W_k_* the inverse variance of the age effect estimate. Similar models are fit for sex by replacing *age_j_* with *sex_j_*. Age at metabolite measured and sex were study-provided by participating studies.

#### Metabolite Imputation

We compared the following missing imputation strategies: no imputation, zero, half-min, median, Quantile Regression for Imputation of Left-Censored data (QRILC)^37^, k-Nearest Neighbors (KNN)^16^, Random Forest (RF)^15^, and Mechanism-Aware Imputation (MAI)^14^. Performance was evaluated based on number of known metQTLs (from prior GWAS in non-TOPMed studies) detected in targeted analysis (and strength of associations) with different missingness imputation strategies, as well as comparisons of number and strength of associations with age and sex, both for known associations and overall (with the assumption that most additional age and sex associations identified are plausible true positives, given the broad impact of both age and sex on the circulating metabolome^11,28,38–44)^.

To assess the downstream effect of imputation methods on genetic association analyses, we first assembled a list of putative metabolite-variant associations based on three large-scale metabolites GWAS studies in studies independent from those included in TOPMed^17,24,45^. After linking metabolites by HMDB, as described above, and filtering by missingness (1-50%) in our datasets, we were able to identify 86 reported metQTLs for 26 metabolites present on the Broad-BIDMC Panel, and 690 metQTLs for 188 metabolites present on the Global Discovery Panel. Treating these associations as “true positives’’, we assessed our ability to replicate these associations for each imputation strategy. Using the GMMAT^46^ pipeline on BioData Catalyst, as described below, we tested for variant-metabolite associations. After imputing metabolite levels and inverse-normalizing values within study sites, we pooled datasets within each platform and tested for genetic associations with adjustment for age, sex, and study (where applicable). To correct for sample relatedness and population structure, we used sparse genetic relationship matrices and the first 11 ancestry principal components. After performing association testing for each imputation method within both metabolomics platforms, we assessed imputation strategies based on the number of literature-reported metQTLs identified as Bonferroni-significant and with the previously reported direction of effect.

### Detection of Metabolite QTLs

Only the metabolites measured in at least two studies and missing rates <50% in each study were included in the following QTL analysis. A total of 1,135 metabolites from 16,359 participants across eight studies were included. The sample size of each study is shown in Supplementary Table 1. The sample size of female and male participants in each study is roughly balanced, except for WHI, which only has female participants. The presence of ancestry diversity in participating studies based on similarity to reference panels has been previously described^47^.

#### Whole Genome Sequencing

TOPMed freeze 10 WGS data was utilized throughout our analyses. Joint calling was performed across all freeze 10 studies by the TOPMed Informatics Research Center (IRC). Detailed calling, processing, and quality control were previously described^47^. In brief, variant calling was completed using the GotCloud pipeline. Variant quality control was performed by calculating Mendelian consistency scores and by applying a support vector machine (SVM) classifier trained on known variant sites and Mendelian inconsistent variants. Sample QC measures included: concordance between annotated and inferred genetic sex, concordance between prior array genotype data and TOPMed WGS data, and pedigree checks. Further details regarding data processing, and quality control are described on the TOPMed website (https://topmed.nhlbi.nih.gov/topmed-whole-genome-sequencing-methods-freeze-9) and in a common document accompanying each TOPMed study’s dbGaP accession. Variants located on all autosomes and the X chromosome were included in both sex-combined and sex-stratified analyses. In this study, we assigned genotypic codes for chromosome X based on the chromosomal region. In non-PAR (pseudoautosomal regions), females were coded as 0, 1, or 2, while males were coded as 0 or 2. In PAR, both females and males were coded as 0, 1, or 2. Variants with minor allele frequency (MAF) less than 0.5% in the overall sample were removed, resulting in 16,518,857 variants on all autosomes and the X chromosome included in the following association analyses.

#### Single variant tests on the pooled TOPMed data

For sex-pooled analysis, a total of 16,359 individuals were analyzed, including 8,855 females and 7,500 males. Missing values on metabolites were imputed using half the minimum values and inverse normal transformations were performed by each study. We included age, sex, study center, and top eleven ancestry principal components as fixed effects covariates^6^. Study_center is the study that the individual belongs to, as well as the recruitment center for relevant studies. The sparse kinship matrix was used as a random effect to account for sample relatedness. A two-stage procedure for rank normalization was applied in genotype-metabolite association analysis^48^. For each of the 1,135 metabolites, we rank-normalized the residuals after regressing the metabolites on fixed-effects covariates and then used them as the outcome in downstream null model fitting and association tests, which was performed by GMMAT pipeline on BioData Catalyst (BDC). We applied a conservative Bonferroni-correction to define genome-wide significance at P-value < 5.0 × 10^−8^/1,135 4.4 x 10^−1^^1^ , accounting for 1,135 analyzed metabolites.

#### Conditional analysis on single variant-metabolite associations

Across the analyzed genome, we defined metabolite-associated genetic loci (i.e., genomic windows) as those containing all statistically significant variants within 500 kb from each other. To account for linkage disequilibrium, we added 500 kbp up- and downstream as a buffer to the genetic loci. If any of these loci overlapped, we merged them into a single locus. In this way, we identified 248 loci, containing 1,248 locus-metabolite pairs.

To identify independent leading variants for each of the locus-metabolite pairs, we conducted conditional analysis by GMMAT. For each locus-metabolites pair, the rank-normalized residuals after regressing the metabolite on fixed-effects covariates from single variant tests were used as the outcome in the downstream null model fitting. The genotypes of those identified top associated variants from each locus together with the fixed-effects covariates were used as covariates in the null model fitting. This conditioning process started with the top associated variant, across the locus followed by a stepwise procedure of selecting additional variants, one by one, according to their conditional *P* values. At the *t-*th step, we calculated the conditional *P* values of all the remaining SNPs in the locus conditional on the SNPs that have been selected in the previous steps. We then selected the SNP with a minimum conditional *P* value that was less than the cutoff *P* value of 5×10^-8^ and added it to the covariates in the next step. We repeated this process until no SNPs could be selected for the model. The variants selected in each step in the conditional analysis were considered conditionally independent.

#### Annotation of known and novel metQTL findings

To access which of the independent variant-metabolite associations were novel, we compiled results from the Metabolomic GWAS Server^1^, TwinsUK Study^22^, GWAS Catalog, GRASP Search, and previous reports from our group^6,20^ and performed a manual search through published papers to detect known loci that overlap with our findings. The significant locus-metabolite pair we identified was classified as known if any variant from a locus-metabolite pair was previously associated with any of the metabolites in its sub-pathway. Otherwise, the locus-metabolite pair was considered novel.

#### Replication of novel metQTLs

We performed replication analysis using metQTLs previously reported among 16,359 multi-ethnic participants from five studies with WGS, including the Atherosclerosis Risk in Communities study (ARIC), Hispanic Community Health Study/Study of Latinos (HCHS/SOL), Cardiovascular Health Study (CHS), FHS and MESA^6^. Of note, a small proportion, 996 out of 16,539 participants (6%), overlapped with the samples from the current analysis. Out of the 108 novel metQTLs, 82 pairs were available for replication in Feofanova et al.^6^. The novel pairs were replicated with either the reported variant or a variant within 500 kb regions for the same metabolite having *P* < 6.10 × 10^−4^. Sixty-six novel pairs were defined to be significantly replicated.

#### Categorization of novel metQTLs based on PhenoScanner

We further used PhenoScanner2 to determine whether the variants from novel metQTLs were also associated with other traits/diseases to improve the function annotation. We got the 135 independent loci (from the 108 novel locus-metabolite associated pairs; Figure 1) and generated the input list by combining all significant variants associated with corresponding metabolites from each independent loci, leading to a total of 6,500 variants for PhenoScanner queries. The scanning was performed on database version 2 (till 12 Dec 2023) with categories setting with “*mQTL*” and “*GWAS*” respectively. We further extracted the “*Disease*’’ related queries from the “*GWAS*” category by matching the “disease” string within traits. The queries from multiple studies were merged as one unique search.

**Figure 1.**
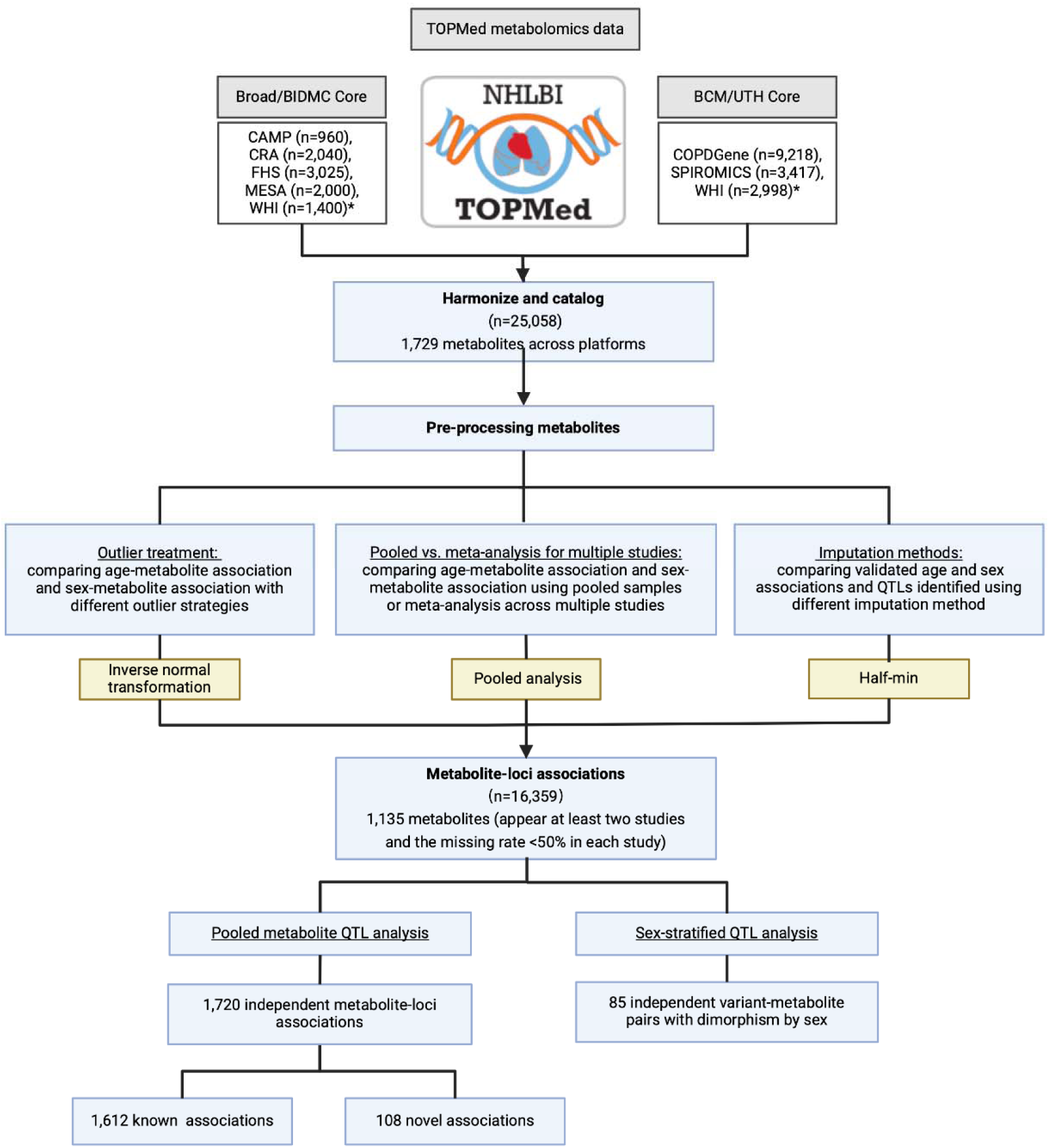
Workflow for comparison of outlier, imputation, and QTL analysis methods. Created with BioRender.com.

### Sex-stratified data analysis

For sex-stratified analysis, the single variant test was conducted on female individuals (8,855 individuals) and male individuals (7,500 individuals) separately following the above analytical steps. Biological sex was defined as “male” for study individuals with sex chromosomes XY and “female” for those with sex chromosomes XX; we acknowledge that such chromosomal make-up data should not be equated with self-identified gender, which was not consistently examined across included studies. We identified the sex-dimorphic associations using the following steps. We first selected the variants that were statistically significant for either females or males, i.e., *P_female_* ≤ 4.4 × 10^−11^ or *P_male_* ≤ 4.4 × 10^−11^. Here, we applied a stringent Bonferroni correction to select female- or male-specific significant associations, accounting for 1,135 analyzed metabolites. Significant variants located within 500 kb of each other were merged into a single locus. To account for linkage disequilibrium, we added 500 kb up- and downstream as a buffer to the genetic loci. If any of these loci overlapped, we merged them into a single locus. In total, 184 independent clumped loci were identified. We then tested if the genetic effects were different between females and males and computed P-values (*P_diff_*). Sex-dimorphic associations were defined with *P_diff_* smaller than the Bonferroni corrected significance level 0.05/184 = 2.7 x 10^−4.^

For each loci-metabolite pair, we conducted the conditional analysis using GMMAT to identify the independently associated variants following the aforementioned steps. The associations of the independent variants that were both statistically significant in the sex-stratified analysis (*P_female_* ≤ 5.0 × 10^−8^ or *P_male_* ≤ 5.0 × 10^−8^) and *P_diff_* ≤ 2.7 × 10^−4^ were considered conditionally independent, sex-dimorphic associations.

## Results

### Catalog of circulating metabolites

We collected metabolite data from 25,058 participants from case-control and population-based studies, including 15,648 participants from three studies profiled on the Global Discovery Panel (i.e., WHI, SPIROMICS, COPDGene), and 9,426 participants from five studies profiled on the Broad-BIDMC panel (i.e., WHI, FHS, CAMP, CRA, and MESA). Metabolites were considered the same if they had the same chemical structure from different platforms. They were ranked normalized in each study separately before they were combined. Across the eight studies, we identified 1,729 named metabolites, which are summarized in Figure 2. There were 366 metabolites identified on both the Broad-BIDMC and the Global Discovery Panels. RefMet annotations showed that named metabolites mostly belonged to organic acids (14%), fatty acyls (21%), glycerolipids (11%), glycerophospholipids (7%), sterol lipids (7%), and nucleic acids (4%). Table 1 summarizes the demographic characteristics of each study with available plasma metabolites funded through TOPMed and for WGS analysis. Overall, samples with metabolites profiled by Broad-BIDMC from WHI had the oldest mean age of 80.0 years with standard deviations (SD) being 7.5, while samples from CAMP had the youngest mean age of 13.0 years with SD being 2.2. All samples with metabolites profiled by the Global Discovery Panel had similar mean ages, ranging from 63.2 years to 66.3 years. Additional information about studies is described in Supplementary Materials. The catalog is provided in Supplementary Table 1, including the HMDB ID, metabolite names based on the latest Metabolon metabolite library and Broad metabolite library, RefMet name, and annotations provided by Metabolomics Workbench.

**Figure 2.**
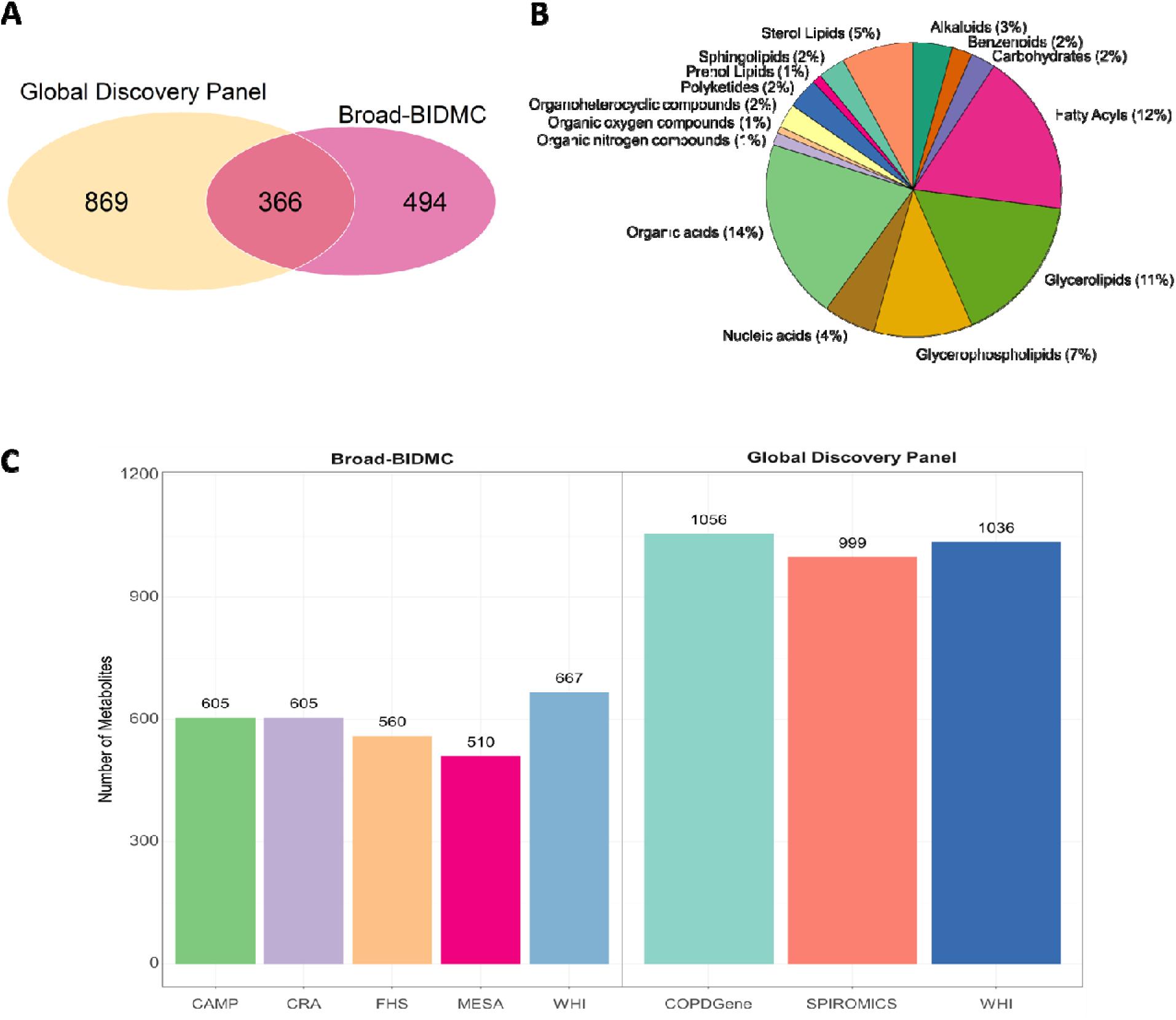
Named metabolites identified from eight studies across two metabolomic cores, th Broad-BIDIMC panel and the Global Discovery Panel. A) Venn diagram of named metabolites measured between platforms; B) Refmet super classes that the named metabolites belong to; and C) the number of named metabolites measured by each study.

**Table 1.** Study samples with available plasma metabolite data from TOPMed. The Overall rows showed study samples with metabolite data available, and the WGS rows showed study samples also with meta-data (i.e., age and sex) and WGS data available, which were included in subsequent analyses.

| Study Name | Study Abbreviation | dbGaP ID | Study Samples | Sample size | Female (%) | Age (mean ± sd) |
| --- | --- | --- | --- | --- | --- | --- |
| Broad-BIDMC Panel |  |  |  |  |  |  |
| The Childhood Asthma Management Program | CAMP | phs001726 | Overall | 960 | 350 (36) | 12.96 (2.15) |
|  |  |  | WGS | 787 | 315 (40) | 13.01 (2.14) |
| The Genetic Epidemiology of Asthma in Costa Rica | CRA | phs000988 | Overall | 2,040 | 918 (45) | 16.38 (14.40) |
|  |  |  | WGS | 1,758 | 806 (46) | 16.78 (14.67) |
| Framingham Heart Study | FHS | phs000974 | Overall | 3,025 | 1,629 (54) | 61.74 (15.15) |
|  |  |  | WGS | 3,021 | 1,626 (54) | 61.76 (15.15) |
| Multi-Ethnic Study of Atherosclerosis * | MESA | phs001416 | Overall | 2,000 | 1,064 (53) | 64.98 (10.72) |
|  |  |  | WGS | 998 | 530 (53) | 60.22 (9.68) |
| Women’s Health Initiative | WHI | phs001237 | Overall | 1,400 | 1,400 (100) | 80.04 (7.54) |
|  |  |  | WGS | 1,024 | 1,024 (100) | 81.69 (6.33) |
| Global Discovery Panel |  |  |  |  |  |  |
| Genetic Epidemiology of COPD | COPDGen e | phs000951<br>phs000946 | Overall | 9,218 | 4,431 (50) | 65.93<br>(9.04) |
|  |  |  | WGS | 6,302 | 3,121 (50) | 64.70<br>(8.86) |
| SubPopulations and Intermediate Outcome Measures in COPD Study | SPIROMICS | phs001927 | Overall | 3,417 | 1,602 (47) | 63.23<br>(9.10) |
|  |  |  | WGS | 1,924 | 888 (46) | 63.32<br>(9.13) |
| Women's Health Initiative | WHI | phs001237 | Overall | 2,998 | 2,998 (100) | 66.33<br>(9.68) |
|  |  |  | WGS | 545 | 545 (100) | 63.65<br>(7.61) |

\*The overall sample size in MESA included participants from visit 1 and visit 5. Only samples from visit 1 were included for WGS analysis.

### Comparison of outlier strategies

For the comparison of outlier and imputation methods, we used pooled analyses to asses associations with age or sex, and metQTLs. However, all outlier handling strategies and missing data imputation were done within studies. To compare the outlier methods in the pooled data, we used complete-case analysis (per-metabolite), by removing missing values (distribution of missingness by study summarized in Supplementary Table 2). For this analysis, we removed any related individuals and removed any metabolites that were only measured in one study. Figure 3A-C compares the parameter estimates of age using 6SD, INT, and winsorization outlier management methods. The same comparison for metabolite-sex associations is provided in Supplementary Figure 1. Agreement of significance of the parameter estimates is indicated by color, using Bonferroni adjusted p-values. All three outlier strategies have very similar estimates for age and sex (minimum correlation > 0.999). Likewise, test statistics for age-metabolite and sex-metabolite associations are highly concordant between outlier strategies (Supplementary Figure 2, Supplementary Table 3). Due to the advantages of normalizing the data, notably the inclusion of extreme values which may be biologically plausible as opposed to dropping subjects altogether, and favorable statistical properties for parametric modeling methods, we continued with INT for the outlier management.

**Figure 3.**
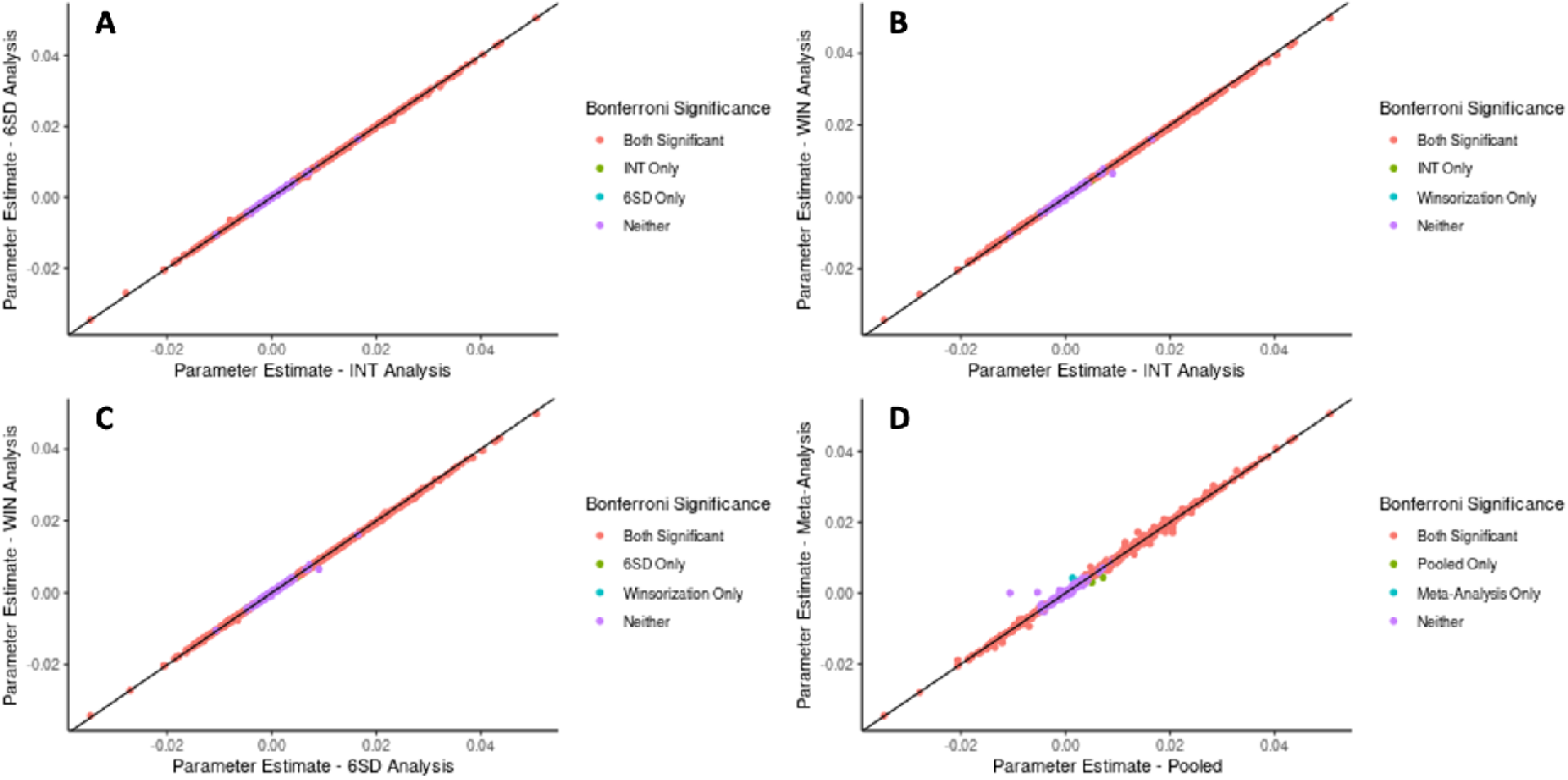
Comparison of age-metabolite associations for outlier handling and data pooling strategies. Parameter estimates for age-metabolite associations are compared for the three outlier handling strategies (A-C) and the two data integration strategies (D).

### Comparison of pooled and meta-analysis

Using INT transformation for outlier management and complete-case analysis, we compared pooled versus study-stratified meta-analysis for age and sex. We note that within-cohort INT would also help address cohort differences in scaling for these semi-quantitative metabolomics measures. As in the previous analysis, we removed related individuals and metabolites measured in only one study. Figure 3D summarizes the effect estimates in the pooled analysis versus the meta-analysis for age for each metabolite and Supplementary Figure 1 summarizes the results for sex. Agreement of significance of the parameter estimates is indicated by color, using Bonferroni adjusted *P* values. In both metabolite age and sex analyses, the parameter estimates and significance have a high agreement (correlation 0.998 and > 0.999 for age and sex, respectively). Full association results are reported in Supplementary Table 4. Our results demonstrate that in most circumstances, pooled and meta-analysis, after inverse normalization by study, give similar results. Our results recapitulate many previously reported associations with age and sex, such as the association of increased leucine (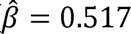, *P* = 4.83 × 10^−202^), carnitine (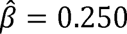, *P* = 812 × 10^−48^), and nicotinamide (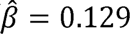, *P* = 6.31 × 10^−15^) with age^49,50^. We also find higher levels of glycine in females (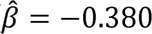, *P* = 4.41 × 10^−109^) and higher levels of creatinine in males (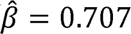, *P* = 1.00 × 10^−323^)^51^.

### Comparison of metabolite imputation strategies

Results for the number and percent of Bonferroni-corrected significant metQTLs detected for each imputation method, within bins of metabolite missingness, are reported in Table 2. Overall, a maximum of 53/85 metQTLs were validated in the Broad-BIDMC panel for the best-performing imputation methods (k-nearest neighbors (KNN), no imputation). In the Global Discovery Panel, a maximum of 512/690 QTLs were validated in the best-performing method (median), a 2.6% improvement over no imputation (499 metQTLs). Of particular interest is imputation quality for metabolites with the highest degree of missingness. Among metabolites with 15-50% missingness, we observed the highest number of QTL-metabolite associations for single value imputation and left-imputation methods (zero, half-min, median, Quantile Regression for Imputation of Left-Censored data (QRILC)^37^ in the Metabolon dataset. However, the total number of significant associations was largely consistent between imputation methods. The distribution of test statistics for metQTL associations is provided in Supplementary Figure 3. A pairwise comparison of these test statistics between each imputation method reveals very similar test statistics for methods that do not assume missing not at random (MNAR) (Random Forest (RF)^15^, Mechanism-Aware Imputation (MAI)^52^, KNN), and similar test statistics for left imputation methods (zero, half-min, QRILC). Full association results are reported in Supplementary Table 5.

**Table 2.** Comparison of quantitative trait loci results by missingness strategy. In the Broad-BIDMC Panel, 85 and 690 literature metQTLs were selected to validate the imputation methods for the Broad-BIDMC Panel and the Global Discovery Panel. The number and percent of Bonferroni-corrected significant metQTLs detected for each imputation method, within bins of metabolite missingness are reported.

| Missingness |  |  |  |  |  |
| --- | --- | --- | --- | --- | --- |
| Platform | Imputation | All | 0-5% | 5-15% | 15-50% |
| Broad-BIDMC Panel | None | 53 (62.4%) | 40 (72.7%) | 7 (30.4%) | 6 (85.7 %) |
|  | Zero | 52 (61.2%) | 38 (69.1%) | 9 (39.1%) | 5 (71.4%) |
|  | Half-min | 52 (61.2%) | 38 (69.1%) | 9 (39.1%) | 5 (71.4%) |
|  | Median | 52 (61.2%) | 38 (69.1%) | 7 (30.4%) | 7 (100%) |
|  | QRILC | 50 (58.9%) | 38 (69.1%) | 9 (39.1%) | 3 (42.9%) |
|  | KNN | 53 (62.4%) | 38 (69.1%) | 9 (39.1%) | 6 (85.7 %) |
|  | RF | 51 (60%) | 38 (69.1%) | 6 (26.8%) | 7 (100%) |
|  | MAI | 52 (61.2%) | 39 (70.9%) | 6 (26.8%) | 7 (100%) |
| Global Discovery Panel | None | 499 (72.3%) | 270 (72.4%) | 180 (75.6%) | 49 (62.0%) |
|  | Zero | 508 (73.6%) | 276 (74.0%) | 182 (76.5%) | 50 (63.3%) |
|  | Half-min | 508 (73.6%) | 276 (74.0%) | 182 (76.5%) | 50 (63.3%) |
|  | Median | 512 (74.2%) | 276 (74.0%) | 183 (76.9%) | 53 (67.1%) |
|  | QRILC | 509 (73.8%) | 276 (74.0%) | 183 (76.9%) | 50 (63.3%) |
|  | KNN | 500 (72.4%) | 271 (72.7%) | 178 (74.8%) | 51 (64.6%) |
|  | RF | 503 (72.9%) | 274 (73.5%) | 179 (75.2%) | 50 (63.3%) |
|  | MAI | 496 (71.9%) | 273 (73.2%) | 176 (73.9%) | 47 (59.5%) |

A comparison of test statistics for imputed metabolite-age and imputed metabolite-sex associations is reported in Supplementary Figure 4. While the distributions per-imputation method is highly similar overall, we observe greater magnitude in the test statistics for left-imputation methods (zero, half-min, QRILC) relative to methods that do not assume MNAR (RF, MAI, KNN). Full association results are reported in Supplementary Table 6.

### Pooled metQTL scanning

We performed a pooled metQTL scanning in our studies, analyzing a total of 16,518,857 variants (MAF>0.5%) on 22 autosomal chromosomes and the X chromosome for associations with 1,135 metabolites. A total of 147,166 single variant-metabolite associations were identified to be significant (*P* ≤ 4.4 × 10^−11^), including 51,866 unique variants covering 667 metabolites. These associations were further grouped into 1,775 conditionally independent locus-metabolite pairs (993 loci, 667 metabolites; Methods and Supplementary Figure 5).

To identify novel associations from these 1,775 conditionally independent locus-metabolite pairs, we further excluded 55 association pairs with unannotated metabolite pathways. Among the remaining 1,720 pairs,1,612 pairs were previously reported or already detected in available public databases (Supplementary Table 7) and 108 locus-metabolite pairs were novel, including 77 loci and 81 metabolites (Supplementary Table 8). Among 108 independent novel pairs, 82 pairs were available for replication, and 80% of loci-trait pairs (66/82) (including 51 unique loci and 51 metabolites) were successfully replicated (*P* ≤ 6.10 × 10^−4^; Supplementary Figure 5).

We then annotated these novel loci with their closest genes. Among them (for lead variants and all significant variants within 500 kb), 1% were classified as missense variants, 5% were in 5 prime UTR regions, 46% were in downstream and upstream regions and others were in non-coding regions (Supplementary Figure 6).

The circular plot in Figure 4 shows the genomic location of significant associations. The genes closest to the novel locus-metabolite pairs were labeled. Of the 77 loci from 108 independent novel association pairs, we observed that 20.8% of loci (16/77) were associated with multiple metabolites, either within the same super-pathways or across super-pathways. Given the one-to-many associations between genes and metabolites, we classified these loci into two categories (Figure 4; Supplementary Table 9). The first category includes 26 loci associated with both reported sub-pathways and unreported sub-pathways in our study. For example, for the known metabolic gene *CD36*, we reproduced a known association of rs3211938 with plasmalogen^23^, and discovered novel associations with sphingomyelins and ceramides. Another example is the rs1047891 located in gene *CPS1* that was reported to be associated with 22 known metabolites^1,6,17,19–21,23,45,53,54^. Meanwhile, we discovered novel associations of *CPS1* with branched-chain fatty acids and partially characterized molecules (Supplementary Table 9). For loci in the second category, we only identified novel metabolite associations. For example, the missense variant rs148051057 located in the *RGN* gene was associated with gluconate level.

**Figure 4.**
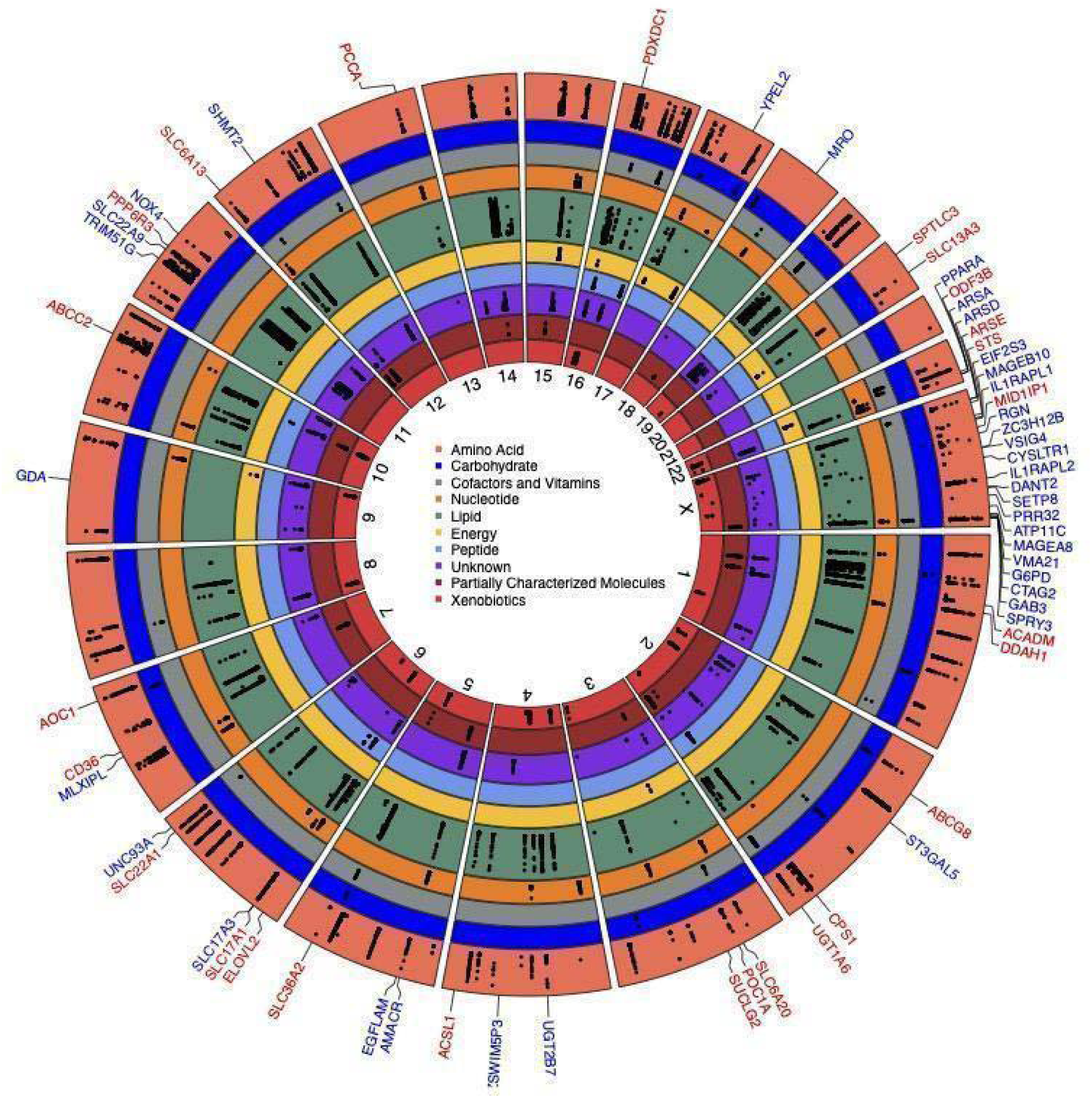
Circular plot illustrating the genomic location of significant associations with metabolites from metabolite pooled analyses. Metabolites were classified into circular bands, and each colored section represented one assigned major pathway class. Significant associations were indicated by black points. The genes closest to the novel locus-metabolites associations were labeled. The genes were classified into two categories: the gene names labeled in red indicated that there were both reported sub-pathways and novel sub-pathways associated with the gene in our study, and the gene names labeled in blue indicated that the gene was only associated with unreported sub-pathways.

Interestingly, we identified 50 novel association pairs from chromosome X. The high percentage of novel pairs (46.3%, 50/108) annotated on chromosome X is in part due to the lack of replication using public databases, as chromosome X variants were not analyzed in many prior studies. The rs112701110 variant near the *MID1IP1* gene was not only associated with the well-known guanidino and acetamido levels^6^, but also had novel associations with leucine, isoleucine, and valine metabolism, and methionine, cysteine, SAM, and taurine metabolism pathway metabolites, etc. A novel association between rs867460874 in the *VSIG4* with long-chain monounsaturated fatty acid was identified (replication *P*-value 3.31 × 10 ). In addition, seven novel loci located in the *GAB3* gene were associated with hemoglobin/porphyrin metabolism and fatty acid metabolism, among which six associations have been replicated (Supplementary Table 8).

Prior work suggested potential links between metQTLs and traits/diseases^8,55^, we therefore performed genotype-phenotype scanning to understand their possible mechanisms (Figure 5). The significant loci were not only detected in complex traits such as height, systolic blood pressure, high-density lipoprotein, total cholesterol, but also overlapped in fat-free mass (including whole body fat-free mass, leg-free mass). Regarding the disease category, the top three enriched diseases are malabsorption/coeliac disease, peripheral vascular disease, and coronary artery disease. The metQTLs related to peripheral vascular disease are from one independent locus (with 14 significant variants) associated with diacylglycerol (DAG) level, including one missense variant rs3812316 located in the *MLXIPL* gene and rs13232120 located in the 3’-UTR of *TBL2* (Supplementary Table 10), indicating shared genetic determinants of DAG-protein kinase C (PKC) pathway and vascular function^56^.

**Figure 5.**
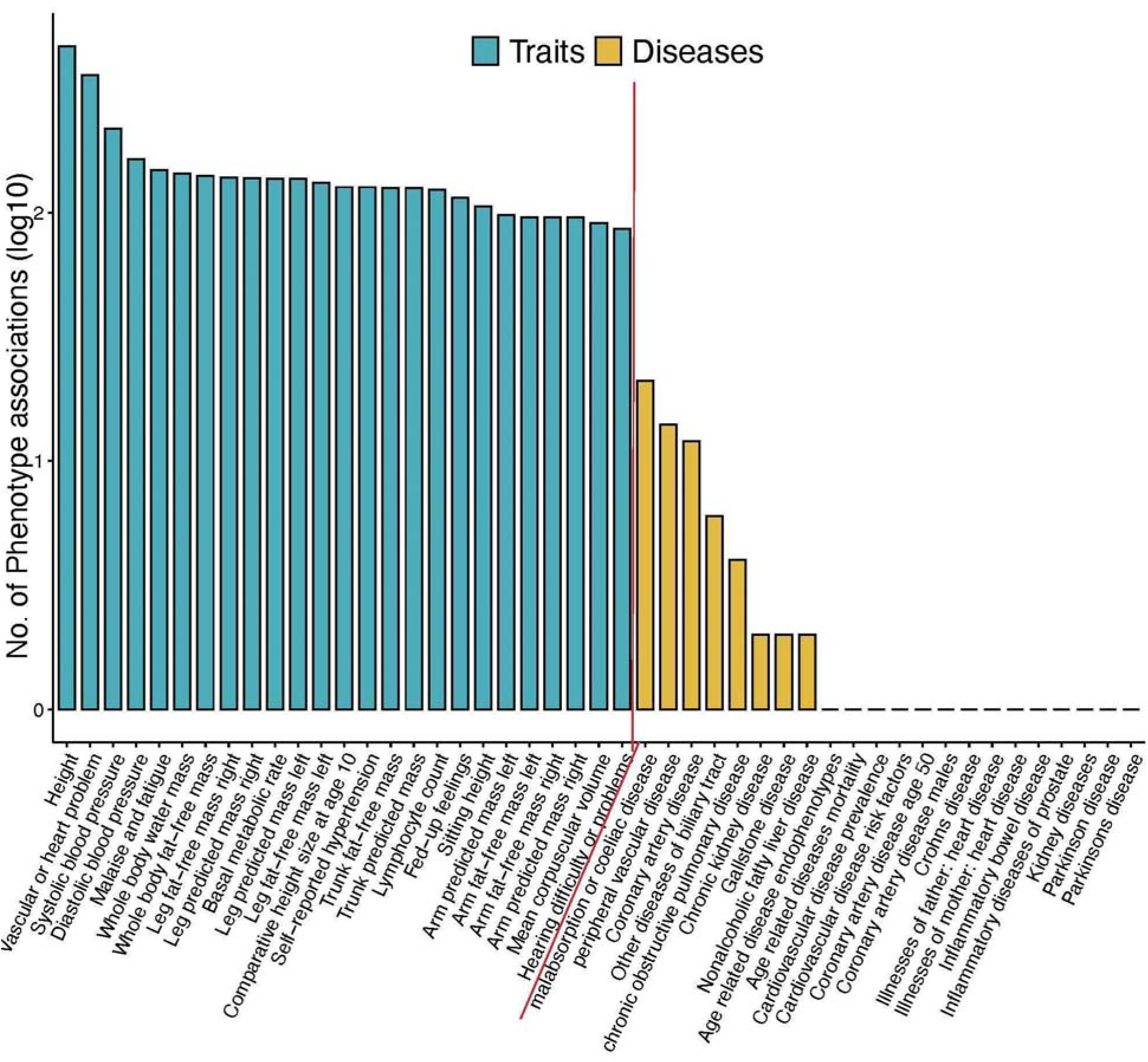
Phenome-wide associations for novel metabolite loci from the pooled analysis. Traits and diseases available in the PhenoScanner (version 2) database were analyzed. The scanning was performed on 7,300 novel significant associated pairs (from 6,500 variants) detected in the pooled analysis (from 108 locus-metabolite pairs). For each category, only the top 25 phenotypes were shown. The y-axis denoted the number of variant hits (with the logarithmic scale) from the PhenoScanner database.

### Sex-stratified metQTL scanning

For the sex-stratified metQTL scanning, 2,453 variant-metabolite associations were identified to show dimorphism by sex, and 85 locus-metabolite pairs were identified to be conditional independent (Supplementary Figure 7; Supplementary Table 11). These independent sex-stratified loci were classified into three categories: 1) significant in females only (44.7%, 33/85); 2) significant in males only (29.4%, 25/85), and 3) significant in both males and female but with different effect sizes (31.8%, 27/85) (Figure 6A-B; Supplementary Figure 8).

**Figure 6.**
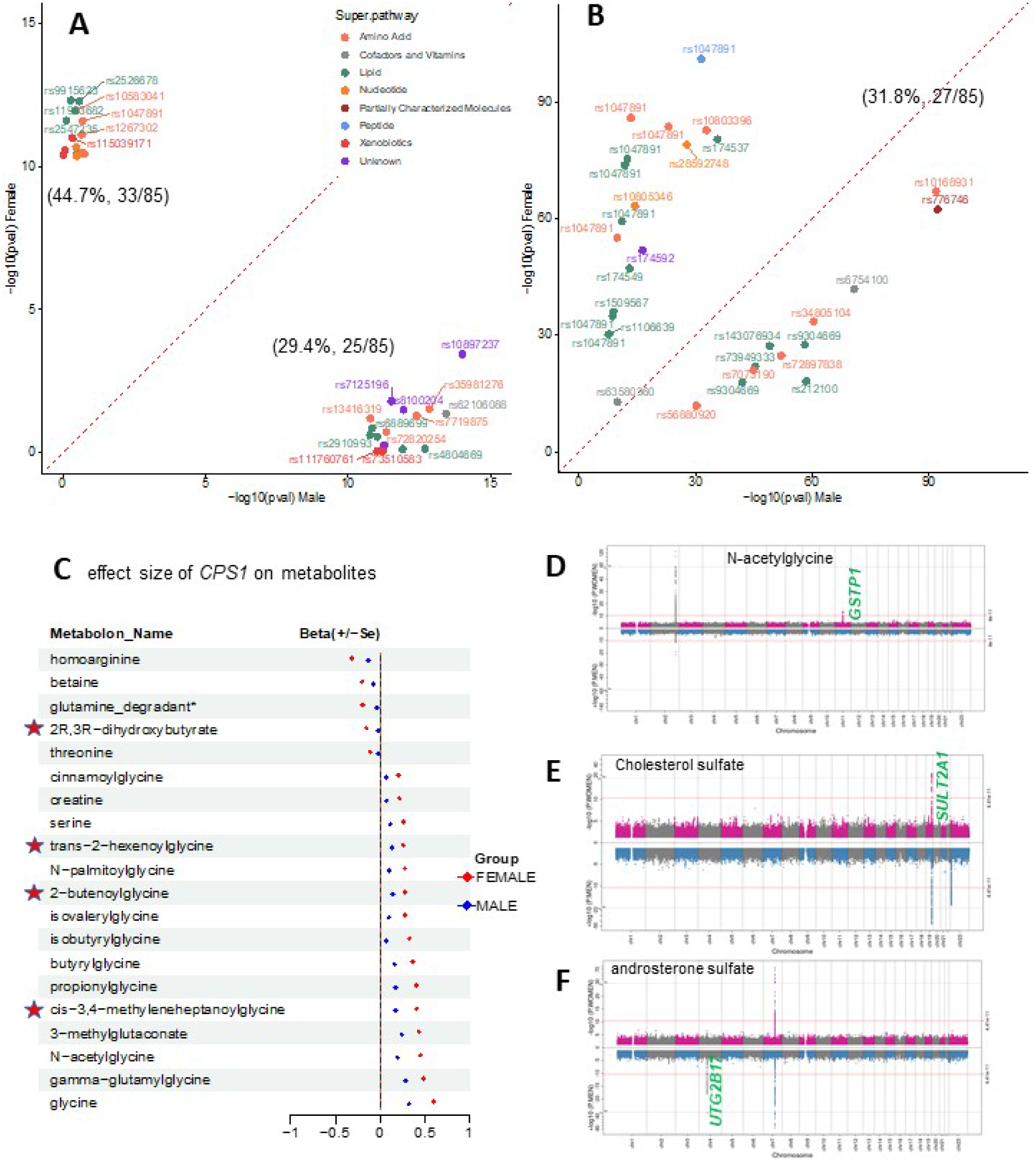
Significant associations identified from sex-stratified analyses. **A**, Scatter plot showing the female- and male-specific association P-values from sex-stratified analysis, the associations are significant for female only (top-left) or male only (bottom-right), **B**, Scatter plot showing significant associations in both males and females but with different effect size level. Each dot denoted one association pair labeled with the rsID and colored by the super pathway of the associated metabolite. Four association pairs with *P* values out of the range (i.e, 0-100) were not shown in **B**), including rs1047891 associated with glycine, N-Acetylglylycine, gamma-Glutamylglycine, and rs7907555 with N2-acetyl,N6-dimethyl lysine. **C**, Sex-stratified per-allele effect size of rs1047891 (in *CPS1*) on metabolite levels. Metabolites not detected in Wittemans et al.^30^ were highlighted with the star symbol. **D-F** Miami-Plot showed available sex-specific association P-value and for trans-2-hexenoylglycine (**D**), cholesterol sulfate (**E**), and androsterone sulfate (**F**) from sex-stratified analyses. The genes mapped by proximity to the sex-specific associations were colored in green.

We observed that rs1047891 located in the *CPS1* gene had the most sex-stratified association pairs, including ten amino acid metabolites, seven lipid metabolites, and one metabolite from partially characterized molecules, peptides, and xenobiotics, separately (Figure 6C). All of them show higher levels in females than males. Importantly, 12 associations were significant in females only, i.e., betaine, creatine, threonine, serine, etc. Among the 20 sex-stratified associations on *CPS1*, 16 associations have been reported. For example, consistent with findings of Wittemans et al.^30^, the effect of the *CPS1* on isobutyrylglycine is nearly 5-fold stronger in females than males (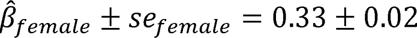, 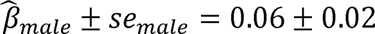, *P_diff_* = 7.49 × 10^−16^). The four metabolites associated with the *CPS1* gene with dimorphism by sex were trans-2-hexenoylglycine, 2-Butenoylglycine, cis-3,4-methyleneheptanoylglycine, and 2R,3R-dihydroxybutyrate with 2-fold, 2-fold, 3-fold, and 7-fold stronger in females than males separately.

We observed an additional 21 association pairs that had significant effects on females only (Supplementary Figure 8A). One example is the association of rs625978 located in *GSTP1* with N-acetylglycine level (Figure 6D; 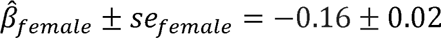, 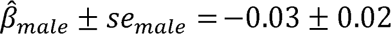, □*P_diff_* < 2.53 × 10^−4^. We also identified associations for well-known metabolic genes such as *FADS2* (with 1,2-dillnoleoyl-GPC), *D2HGDH* (with 2-hydroxyglutarate), *SULT2A1* (with 5alpha-androstan-3beta), *ASMTL* (with gualacol sulfate), and *ACADL* (with nonanoylcarnitine) etc.

25 association pairs only showed significant effects in males (Supplementary Figure 8B). We observed the rs34707604 in *UTG2B17* associated with three metabolites of lipid androgenic steroids, including androsterone sulfate (Figure 6F), 5alpha-androstan-3alpha,17beta-diol monosulfate (1), and androstenediol (3alpha, 17alpha) monosulfate (3). Its effect on androsterone sulfate was nearly 4-fold stronger in males than females (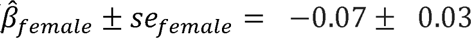, 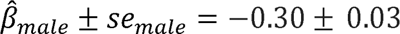, □*P_diff_* < 5.3 × 10^−9^). Similarly, there were male-specific associations for well-known metabolic genes such as *PTPRN2* (with propyl 4-hydroxybenzoate sulfate), *UGT2B17* (with androsterone sulfate), *SUGP1* (with Cer 18:1;O2/24:0), *ABCC1* (with glutarylcarnitine (C5-DC)), *UGT3A1* (with 5alpha-pregnan-3beta).

There were four sex-stratified loci located on the X chromosome. The top significant signal was rs34805104 located in *TMLHE-SPRY3* associated with N6-trimethyllysine level, with the effect size estimate in males was nearly 2-fold of that in females (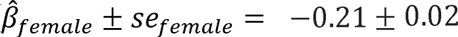, 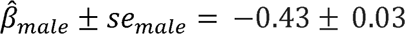, □*P_diff_* < 2.02 × 10^−12^).

## Discussion

In this study, we harmonized and cataloged circulating metabolites from eight TOPMed studies. We demonstrated one set of reasonable strategies for both genetic and epidemiological association analyses within the TOPMed program, concerning methods for handling outliers, missing data, and pooled versus meta-analysis. We expect this analysis framework will prompt increasing use of the valuable TOPMed metabolomics resources. We also conducted a WGS study to detect genetic association with 1,135 circulating metabolites. Among 108 novel locus-metabolite pairs, we detected not only novel associations for known metabolic genes, but also novel genes that have putative roles in metabolic regulation based on prior literature. The sex-stratified analysis also provided a systematic, large-scale examination of sex differences in the genetic regulation of the circulating metabolites.

Metabolite overlap across five metabolomic platforms (the Global Discovery Panel, the Broad-BIDMC Panel, Nightingale Health, Biocrates, and West Coast Metabolomics) has previously been assessed by COMETS^57^. Between the two major metabolite platforms, there were 1,155 metabolites measured by the Global Discovery Panel, 350 metabolites measured by the Broad-BIDMC Panel, and 111 overlapping metabolites identified using names and identifiers from the Human Metabolome Database, Pubchem, and Chemspider. In our current assessment of metabolite overlap across eight TOPMed studies, we identified 1,729 total metabolites with known identities and 366 overlapping metabolites between the Global Discovery Panel and Broad-BIDMC Panel. The number of identified metabolites overlapping between these two popular metabolite platforms was more than tripled with the latest metabolite libraries used in TOPMed. Our current work has harmonized additional metabolites that previously may not have been measured or had a known identity, expanding the scope of metabolomic research.

With technological advances in liquid chromatography combined with mass spectrometry (LC-MS), the collection of metabolomic data has increased. However, LC-MS generated data is missingness. This missingness could be true missing (i.e., the metabolite is absent from the biological samples, the true concentration of zero), missing due to technical factors (such as co-elution), or could be the level is below the detection limit. There are two main approaches to handling missingness, (1) to impute the data and use statistical methods that require complete data or (2) to use statistical methods that can account for missingness without imputation. In the first approach, previous studies have compared different methods to handle or impute the missingness. Kokla et al.^14^ simulated different missingness mechanisms, i.e., missing completely at random (MCAR), missing not at random (MNAR), and missing at random (MAR), and compared imputation methods including zero, mean, minimum value, half minimum, singular value decomposition, probabilistic principal component analysis, Bayesian principal component analysis, random forest, and k-nearest neighbors. In the MCAR and MAR scenarios, RF and KNN performed the best. For our analysis, assuming that previously reported QTLs from GWAS studies of age and sex in non-TOPMed studies were true positives, we found that half min performed similarly to RF or KNN. In the case where there is high missingness (15-50%), we found that KNN, RF, and MAI have the best performance. MAI is also the recommended method by the Metabolomics Workbench^58^. Other studies also concluded that RF tended to outperform other methods in performance^59^. RF imputation was found to perform well for MAR and MCAR, but poorly when the data was MNAR^37^. For MNAR, they recommended quantile regression for the imputation of left-censored data (QRILC), which we also examined. In the second approach, imputation-free methods have been previously compared with standard imputation methods, and the results indicated that no imputation-free method was consistently better^58^. We find very few meaningful differences across imputation methods for the Broad-BIDMC panel, and some improvements with left-imputation methods for the Global Discovery Panel. There may be differences in the proportion of missing data that is MAR versus MNAR between the two platforms, and we note that there are both many more named metabolites and many more named metabolites with high missingness for the Global Discovery Panel, which may impact our comparisons. In both the Global Discovery Panel and Broad-BIDMC Panel, our results suggest that simpler approaches may provide comparable performance to more complex methods (MAI, kNN, etc). Accounting for the computation burden and technical complexity, we have recommended the use of half minimum value for most TOPMed analyses; this approach was utilized in the genome-wide metabolite QTL associations provided here.

Multi-study and integrative metabolomic analyses are often conducted via a meta-analytic approach^9–12^, where association studies are conducted per study, and effect sizes or *P* values are combined. Meta-analyses can account for study-specific considerations and overcome many privacy issues where data cannot be shared and pooled in one analysis location. However, meta-analyses may also have bias and lower power to detect individual-level interactions^60^. Particularly in the scope of rare-variant analysis, meta-analysis has many challenges. Pooling data has been suggested as an alternative approach to meta-analysis^13^. Pooled data increases the sample size, has greater flexibility to control confounders, and avoids within-study assumptions. Pooling data allows for a better ability to detect rare outcomes efficiently, especially if the outcome is low in individual studies, for example when assessing associations of rare genetic variants. However, the pooling analysis requires the data to be homogenous and more post-pooling cleaning, formatting, and resources for analysis. Feofanova et al.^6^ pooled five studies, including MESA, and metabolomic platforms with up to 1666 circulating metabolites, and performed whole-genome sequencing association analysis. They successfully identified 75 novel and replicated metabolite-genetic locus associations. The pooled data identified novel rare-variant gene pairs, while effectively controlling for genomic inflation. Additionally, they were able to extend metabolite genetic association discovery to multi-ethnic populations^61^. We similarly show the feasibility and power of a study-pooled analytic approach with appropriate outlier management via inverse normalization by study before analysis.

In our pooled association analyses, 93.7% (1612/1720) of associations overlapped with findings from previously reported studies, demonstrating the high reliability of metQTLs. For example, with association scanning on glycine, we identified four significant loci located in genes *CPS1*, *ALDH1L1*, *PSPH*, and *GCSH* (Supplementary Figure 9). Remarkably, all these four genes are essentially involved in the glycine metabolism pathway^30^. We were also able to reproduce well-known signals from the sex-stratified analysis. For example, most metabolites showing female-specific associations for rs1047891 located in *CPS1* in our study have been detected in a previous study^30^.

Among the 66 novel and replicated pairs from the pooled analysis, nearly half of these detected involved the X chromosome, which has not been included in many prior genetic analyses. For example, five novel and replicated pairs were identified between the *GAB3* locus and bilirubin, biliverdin, cerotoylcarnitine (C26), palmitoylcarnitine (C16), and stearoylcarnitine (C18). Cerotoylcarnitine (C26), palmitoylcarnitine (C16), and stearoylcarnitine (C18) are acylcarnitines involved in fatty acid metabolism. The *GAB3* gene encodes the protein that binds to the growth factor receptor-bound protein 2 (*GRB2*). *GRB2* plays a central role in signaling by receptor protein-tyrosine kinases (RTKs)^62,63^, which respond to the FGF receptor family to affect fatty acid synthesis^64^. Our association scanning indicates the putative role of *GAB3* in controlling lipid metabolism by the mediation of RTK ligands. Another novel association of rs867460874 in the *VSIG4* locus with myristoleate (14:1n5) in the sub-pathway of long-chain monounsaturated fatty acid was identified. *VSIG4*, a B7 family-related protein expressed by resting macrophages, can regulate pyruvate/acetyl-CoA conversion by activating the PI3K/Akt-STAT3 pathway^65^. Since pyruvate and acetyl-CoA are important intermediaries in the conversion of carbohydrates into fatty acids and cholesterol^66^, this may reveal the putative link of the *VSIG4* gene on lipid metabolism.

The influence of sexual dimorphism is gaining mounting interest in human metabolomic research^67,68^. We detected that *GSTP1* only showed female-specific effects on N-acetylglycine level. This gene encodes glutathione S-transferase enzymes and plays an important role in glutathione metabolism and drug metabolism^69^. Polymorphisms in this gene have been reported to be associated with male infertility^70^, supporting the potential sex-stratified role of *GSTP1* on metabolite level. *UGT2B17* only shows male-specific effects on lipid androgenic steroid levels.

This gene encodes an enzyme for glucuronidation of androgens and their metabolites, which is a major source of estrogen^28,71^. A higher concentration of dehydroeplandrosterone sulfate in male participants has been reported^71^, further supporting the potential sex-stratified role of *UTG2B17* on androsterone sulfate level. Furthermore, we detected well-known metabolic genes, such as *CPS1, FADS2, D2HGDH*, and *SUGP1*, showing sex-stratified effects on multiple metabolite levels, which requires further investigation. Taken together, our study strongly supports the importance of sex-stratified analysis for studying the underlying mechanisms of sex difference in human metabolomes.

TOPMed has a variety of strengths and weaknesses versus other large-scale efforts to profile the circulating metabolome. TOPMed is a program of many studies, with centralized metabolomic cores, but with varying measured phenotypes and recruitment strategies. COMETS has a very large total sample size (>136,000 samples with metabolomics from >47 studies). However, due to widely varying study designs and laboratory methods and data sharing restrictions, conducting pooled analyses is challenging as compared to the TOPMed program^57^. UK Biobank has generated metabolomics measures in an extremely large sample size, as have other biobanks such as Estonian Biobank and Finnish THL Biobank^72^, on the targeted Nightingale platform. This platform has advantages, particularly in terms of scalability and cost, but covers a narrow slice of the circulating metabolome, mostly from amino acids and lipids^73^. Untargeted metabolomic platforms have been used in TOPMed, as well as the inclusion of many studies with diverse participants from across the life course, which enables the capture of a variety of metabolites from different pathways for a comprehensive assessment of impact from human metabolomes. TOPMed data generation, including longitudinal data generation, is ongoing, and sample sizes in diverse TOPMed studies will continue to accrue. Pooled analysis makes it more feasible for certain types of analysis strategies, such as aggregate tests for rare variants^6^. Our results show a feasible strategy for conducting cross-study analyses for the two different platforms in TOPMed.

Future analysis will also work to integrate both TOPMed generated and study generated metabolomics data to promote the discovery of metQTLs, from both single variant and aggregate rare variant tests. We will also work to incorporate our analytical framework and to extend our harmonization efforts to additional metabolomic samples funded by TOPMed (with metabolomic quantification for >90,000 samples, including longitudinal measurements, across 13 studies currently funded). Our current work has focused on named metabolites; additional insights may be gained by examining unnamed compounds, with genetic analysis potentially providing clues to metabolite identity^23,74,75^. Such work will further elucidate our understanding of the genetic and epidemiological determinants of the circulating metabolome in diverse US populations.

In summary, we here describe the first set of TOPMed generated metabolomics data across two platforms (the Global Discovery Panel and Broad-BIDMC Panel) and eight diverse studies. A metabolite catalog was created and released on the TOPMed website and as supplementary data with this publication. We evaluate the impact on results of multiple common analysis strategies for handling outlier and missing values, as well as comparing pooled versus meta-analysis strategy. We find that within the study inverse normalization and half-minimum imputation for missing data provide reasonable and well-powered results in both pooled and meta-analyses. We utilize these strategies for cross-study metQTL analyses, identifying known and novel loci and validating the high quality of the metabolomics resources generated. TOPMed generated individual-level data (Supplementary Table 12) and summary metQTL data (phs001974) are available for access through dbGaP, providing a resource for the scientific community to integrate metabolomics for understanding common complex diseases.

## Data and code availability

The genotype data and metabolomics data used in this paper are available on request through the dbGaP (CAMP with phs001726, COPDGene with phs000951 and phs001927, CRA with phs000988, FHS with phs000974, MESA with phs001416, SPIROMICS with phs001927, WHI with phs001237). The metabolite catalog is available at https://topmed.nhlbi.nih.gov/omics. The summary statistics are available under dbGaP accession number: phs001974.v5.p1.

## Acknowledgments

Molecular data for the Trans-Omics in Precision Medicine (TOPMed) program was supported by the National Heart, Lung, and Blood Institute (NHLBI). See the TOPMed Omics Support section in Supplementary Materials for study specific omics support information. Core support including centralized genomic read mapping and genotype calling, along with variant quality metrics and filtering were provided by the TOPMed Informatics Research Center (3R01HL-117626-02S1; contract HHSN268201800002I). Core support including phenotype harmonization, data management, sample-identity QC, and general program coordination was provided by the TOPMed Data Coordinating Center (R01HL-120393; U01HL-120393; contract HHSN268201800001I). We gratefully acknowledge the studies and participants who provided biological samples and data for TOPMed. Additional study specific acknowledgments are included in Supplementary Materials.

NF is supported by NIH grants R01MD012765, R01DK117445, and R01HL163972. RSK is supported by NHLBI grant: K01HL146980. LW is supported by NHGRI/NIMHD U54HG013243. The work and BY are in part supported by R01HL168638 and the JLH Foundation.

## Declaration of interests

In the past three years, EKS received grant support from Bayer and Northpond Laboratories. LW provided consulting services to Pupil Bio Inc. and reviewed manuscripts for the Gastroenterology Report, not related to this study, and received an honorarium. LMR is a consultant for the TOPMed Administrative Coordinating Center (through Westat).

## Notes

### Competing Interest Statement

The authors have declared no competing interest.

### Summary of Updates

Add supplemental files, and some polish on manuscript

