## Supplemental Information for "Genetic Architecture and Analysis Practices of Circulating Metabolites in the NHLBI Trans-Omics for Precision Medicine (TOPMed) Program"

### Supplementary Information

#### Cohort Description

**COPDGene** The multi-center, NIH-funded COPDGene (also known as the Genetic Epidemiology of COPD Study) includes a study population of more than 10,000 smokers (~1/3 self-reported non-Hispanic Black and 2/3 non-Hispanic White participants) and has been characterized with a study protocol including pulmonary function tests, chest CT scans, six-minute walk testing, and multiple questionnaires[1]. Five years and ten years after this initial visit a similar protocol was conducted at a follow-up visit. More details are available at http://www.copdgene.org/.

**FHS** The Framingham Heart Study (FHS) is a three-generation, single-site, community-based, ongoing cohort study initiated in 1948 to investigate prospectively the risk factors for CVD[2–4]. It now comprises 3 generations of participants: the original cohort followed since 1948; their Offspring and spouses of the Offspring, followed since 1971; and children from the largest Offspring families enrolled in 2002 (Gen 3). The Original cohort enrolled 5,209 men and women (comprising two-thirds of the adult population in Framingham, MA at that time). Survivors continue to receive biennial examinations. The Offspring cohort comprises 5,124 persons (including 3,514 biological offspring) who have been examined approximately once every 4 years. The Gen 3 cohort contains 4,095 participants.

**MESA** The Multi-Ethnic Study of Atherosclerosis (MESA) is designed to assess subclinical cardiovascular disease and the risk factors that predict cardiovascular disease progression. MESA recruited a diverse, population-based sample of 6,814 asymptomatic men and women aged 45-84 at baseline[5]. In terms of self-reported race and/or ethnicity, 38 percent of the recruited participants were non-Hispanic White, 28 percent Black or African-American, 22 percent Hispanic, and 12 percent Asian, predominantly of Chinese descent. Participants were recruited from six field centers across the United States: Wake Forest University, Columbia University, Johns Hopkins University, University of Minnesota, Northwestern University and the University of California - Los Angeles.

**WHI** The Women's Health Initiative (WHI) is a long-term, prospective, multi-center cohort study that investigates post-menopausal women's health, including strategies to prevent heart disease, breast cancer, colon cancer, and osteoporotic fractures[6]. WHI recruited 161,808 women between 1993 and 1998 at 40 centers across the US (50-79 years at baseline). The study consists of two parts: the WHI Clinical Trial, a randomized clinical trial of hormone therapy, dietary modification, and calcium/Vitamin D supplementation, and the WHI Observational Study, assessing the incidence, risk factors, and interventions related to heart disease, cancer, and osteoporotic fractures. WHI samples sequenced in TOPMed were selected on a case/control basis for stroke and venous thromboembolism.

**CRA** The Genetic Epidemiology of Asthma in Costa Rica (CRA) Cohort recruited 4,245 participants, focusing on children with asthma, their parents, and other pedigree relatives, after screening over 9,180 children enrolled in Costa Rican schools[7]. All probands impacted by asthma completed a protocol including questionnaires, spirometry, methacholine challenge testing (if FEV1 was ≥ 65% of predicted), allergy skin testing, and collection of blood (for plasma, DNA and RNA extraction, and measurement of serum total and allergen-specific IgE) and house dust (for measurement of dust mite/cockroach allergens) samples.

**CAMP** As described previously, the Childhood Asthma Management Program (CAMP) recruited 1,041 children from 5–12 years of age with mild-to-moderate persistent asthma from 1993-95 at eight clinical centers[8]. After initial baseline screening to establish asthma severity, participants were randomized to budesonide dry-powder inhaler (200 μg twice daily), nedocromil metered-dose inhaler (8 mg twice daily), or matching placebos for 4–6 years.

**SPIROMICS** The Subpopulations and Intermediate Outcomes in COPD Study (SPIROMICS) is a multi-center observational study of chronic obstructive pulmonary disease (COPD), with 2,982 participants aged 40-80 at baseline in the main study[9]. Most participants had current or former tobacco use for longer than 20 pack-years; control participants without a history of smoking or lung disease were also recruited. Participants participated in deep phenotyping related to COPD, including spirometry and a high-resolution chest CT scan and repeat questionnaires regarding tobacco use, pulmonary symptoms (COPD Assessment Test score), respiratory exacerbations, use of oral corticosteroids and antibiotics, emergency department (ED) visits and hospitalizations. More information is available at https://www.spiromics.org/.

### Study Specific Acknowledgments

**NHLBI TOPMed: Genetic Epidemiology of COPD Study (COPDGene)**

The COPDGene study (NCT00608764) is supported by grants from the NHLBI (U01HL089897 and U01HL089856), by NIH contract 75N92023D00011. The content is solely the responsibility of the authors and does not necessarily represent the official views of the National Heart, Lung, and Blood Institute or the National Institutes of Health. The COPDGene project is also supported by the COPD Foundation through contributions made to an Industry Advisory Committee that has included AstraZeneca, Bayer Pharmaceuticals, Boehringer-Ingelheim, Genentech, GlaxoSmithKline, Novartis, Pfizer and Sunovion. A full listing of COPDGene investigators can be found at: http://www.copdgene.org/directory

**NHLBI TOPMed: Framingham Heart Study (FHS)**

The Framingham Heart Study (FHS) acknowledges the support of contracts NO1-HC-25195, HHSN268201500001I and 75N92019D00031 from the National Heart, Lung and Blood Institute and grant supplement R01 HL092577-06S1 for this research. We also acknowledge the dedication of the FHS study participants without whom this research would not be possible. Dr. Vasan is supported in part by the Evans Medical Foundation and the Jay and Louis Coffman Endowment from the Department of Medicine, Boston University School of Medicine.

**NHLBI TOPMed: Multi-Ethnic Study of Atherosclerosis (MESA)**

Please contact the MESA Genetics P&P Coordinator for MESA acknowledgements. MESA and the MESA SHARe projects are conducted and supported by the National Heart, Lung, and Blood Institute (NHLBI) in collaboration with MESA investigators. Support for MESA is provided by contracts 75N92020D00001, HHSN268201500003I, N01-HC-95159, 75N92020D00005, N01-HC-95160, 75N92020D00002, N01-HC-95161, 75N92020D00003, N01-HC-95162, 75N92020D00006, N01-HC-95163, 75N92020D00004, N01-HC-95164, 75N92020D00007, N01-HC-95165, N01-HC-95166, N01-HC-95167, N01-HC-95168, N01-HC-95169, UL1-TR-000040, UL1-TR-001079, and UL1-TR-001420, UL1TR001881, DK063491, and R01HL105756. Funding for SHARe genotyping was provided by NHLBI Contract N02-HL-64278. Genotyping was performed at Affymetrix (Santa Clara, California, USA) and the Broad Institute of Harvard and MIT (Boston, Massachusetts, USA) using the Affymetrix Genome-Wide Human SNP Array 6.0. The authors thank the other investigators, the staff, and the participants of the MESA study for their valuable contributions. A full list of participating MESA investigators and institutes can be found at http://www.mesa-nhlbi.org.

**NHLBI TOPMed: SubPopulations and InteRmediate Outcome Measures In COPD Study (SPIROMICS)**

The authors thank the SPIROMICS participants and participating physicians, investigators, study coordinators, and staff for making this research possible. More information about the study and how to access SPIROMICS data is available at www.spiromics.org. The authors would like to acknowledge the University of North Carolina at Chapel Hill BioSpecimen Processing Facility (http://bsp.web.unc.edu/) and Alexis Lab (https://www.med.unc.edu/cemalb/facultyresearch/alexislab/) for sample processing, storage, and sample disbursements.

We would like to acknowledge the following current and former investigators of the SPIROMICS sites and reading centers: Neil E Alexis, MD; Wayne H Anderson, PhD; Mehrdad Arjomandi, MD; Igor Barjaktarevic, MD, PhD; R Graham Barr, MD, DrPH; Patricia Basta, PhD; Lori A Bateman, MS; Christina Bellinger, MD; Surya P Bhatt, MD; Eugene R Bleecker, MD; Richard C Boucher, MD; Russell P Bowler, MD, PhD; Russell G Buhr, MD, PhD; Stephanie A Christenson, MD; Alejandro P Comellas, MD; Christopher B Cooper, MD, PhD; David J Couper, PhD; Gerard J Criner, MD; Ronald G Crystal, MD; Jeffrey L Curtis, MD; Claire M Doerschuk, MD; Mark T Dransfield, MD; M Bradley Drummond, MD; Christine M Freeman, PhD; Craig Galban, PhD; Katherine Gershner, DO; MeiLan K Han, MD, MS; Nadia N Hansel, MD, MPH; Annette T Hastie, PhD; Eric A Hoffman, PhD; Yvonne J Huang, MD; Robert J Kaner, MD; Richard E Kanner, MD; Mehmet Kesimer, PhD; Eric C Kleerup, MD; Jerry A Krishnan, MD, PhD; Wassim W Labaki, MD; Lisa M LaVange, PhD; Stephen C Lazarus, MD; Fernando J Martinez, MD, MS; Merry-Lynn McDonald, PhD; Deborah A Meyers, PhD; Wendy C Moore, MD; John D Newell Jr, MD; Elizabeth C Oelsner, MD, MPH; Jill Ohar, MD; Wanda K O'Neal, PhD; Victor E Ortega, MD, PhD; Robert Paine, III, MD; Laura Paulin, MD, MHS; Stephen P Peters, MD, PhD; Cheryl Pirozzi, MD; Nirupama Putcha, MD, MHS; Sanjeev Raman, MBBS, MD; Stephen I Rennard, MD; Donald P Tashkin, MD; J Michael Wells, MD; Robert A Wise, MD; and Prescott G Woodruff, MD, MPH. The project officers from the Lung Division of the National Heart, Lung, and Blood Institute were Lisa Postow, PhD, and Lisa Viviano, BSN; SPIROMICS was supported by contracts from the

NIH/NHLBI (HHSN268200900013C, HHSN268200900014C, HHSN268200900015C, HHSN268200900016C, HHSN268200900017C, HHSN268200900018C, HHSN268200900019C, HHSN268200900020C), grants from the NIH/NHLBI (U01 HL137880, U24 HL141762, R01 HL182622, and R01 HL144718), and supplemented by contributions made through the Foundation for the NIH and the COPD Foundation from Amgen; AstraZeneca/MedImmune; Bayer; Bellerophon Therapeutics; Boehringer-Ingelheim Pharmaceuticals, Inc.; Chiesi Farmaceutici S.p.A.; Forest Research Institute, Inc.; Genentech; GlaxoSmithKline; Grifols Therapeutics, Inc.; Ikaria, Inc.; MGC Diagnostics; Novartis Pharmaceuticals Corporation; Nycomed GmbH; Polarean; ProterixBio; Regeneron Pharmaceuticals, Inc.; Sanofi; Sunovion; Takeda Pharmaceutical Company; and Theravance Biopharma and Mylan/Viatris.

**NHLBI TOPMed: Women's Health Initiative (WHI)**

The WHI program is funded by the National Heart, Lung, and Blood Institute, National Institutes of Health, U.S. Department of Health and Human Services through contracts 75N92021D00001, 75N92021D00002, 75N92021D00003, 75N92021D00004, 75N92021D00005.

### Supplementary Figures

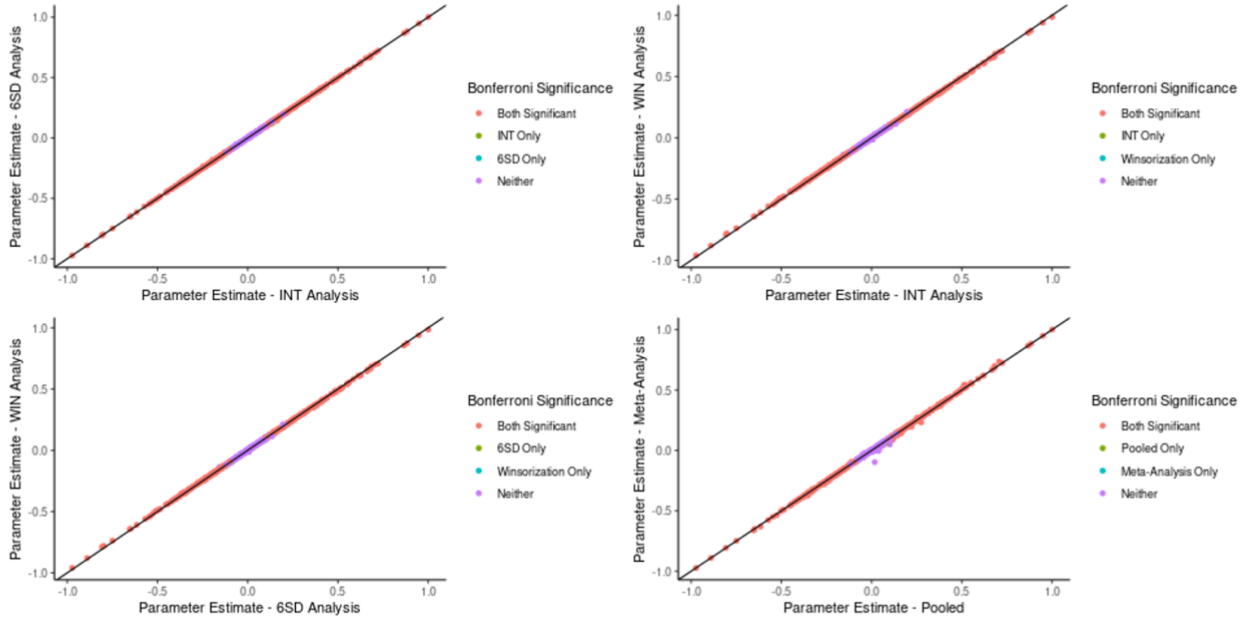


**Supplementary Figure 1. Comparison of sex-metabolite associations for outlier handling and data pooling strategies.** Parameter estimates for sex-metabolite associations are compared for the three outlier handling strategies (A-C) and for the two data integration strategies (D).

**(Age-metabolite Z-scores)**

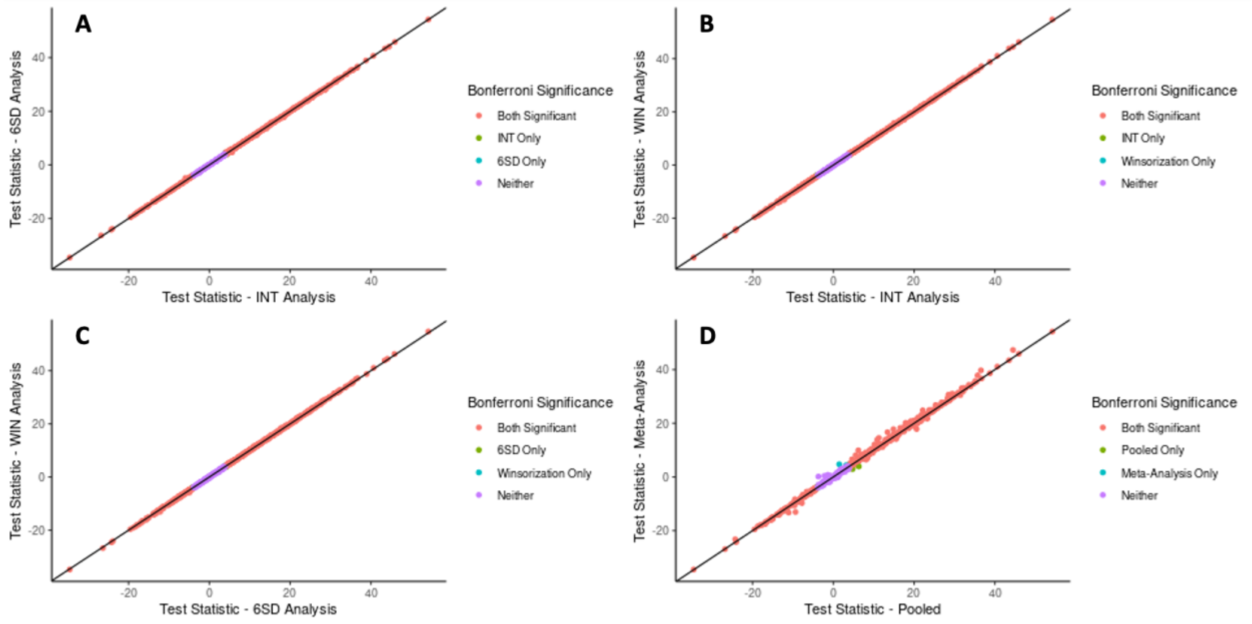


**(Sex-metabolite Z-scores)**

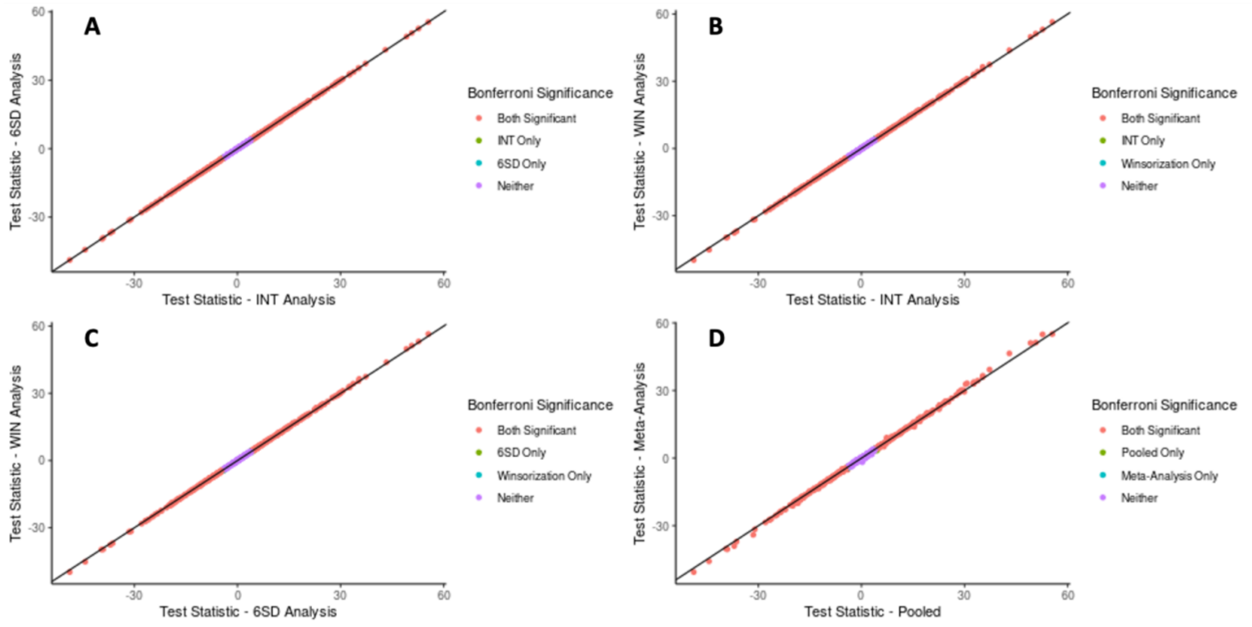


**Supplementary Figure 2. Comparison of test statistics for outlier handling and data pooling strategies.** The upper panel shows the result for age-metabolite association, the lower panel shows the result for sex-metabolite associations. In each panel, (A-C) is the comparison of test statistics for the three outlier handling strategies and (D) is the comparison of test statistics for the two data integration strategies.

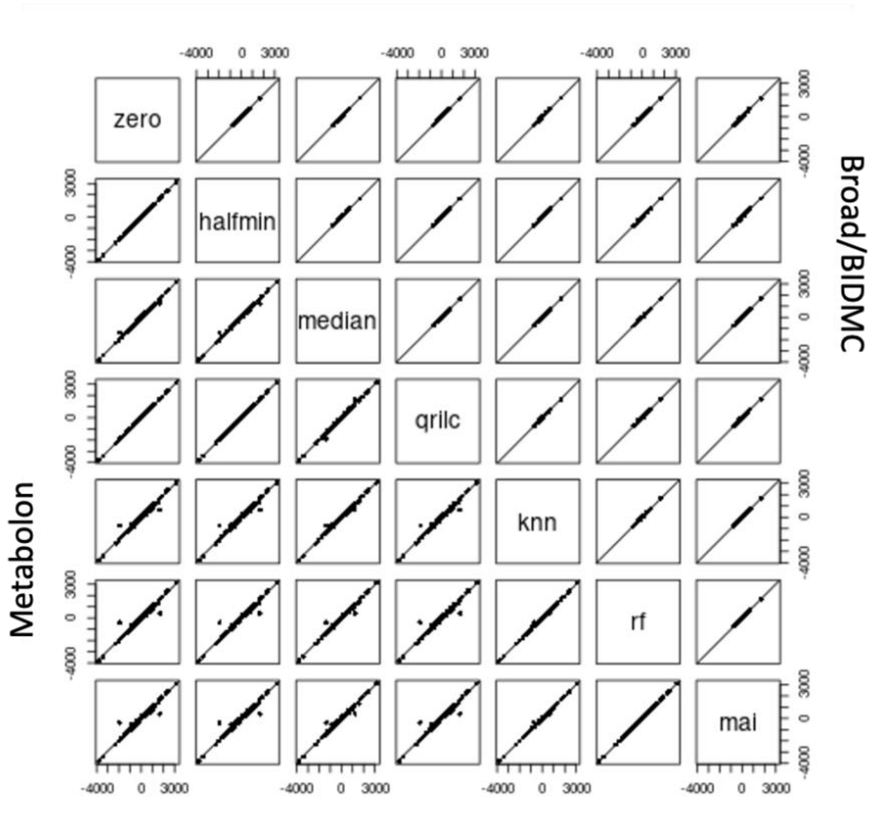


**Supplementary Figure 3. The distribution of test statistics for metQTL associations for different imputation strategies.**

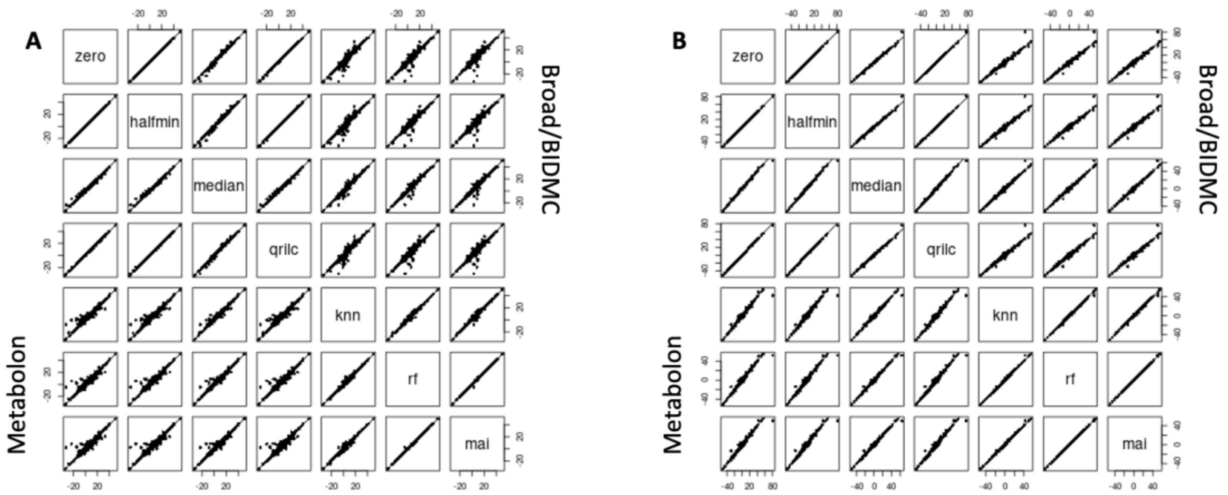


**Supplementary Figure 4. Comparison of test statistics for imputed metabolite-age (A) and imputed metabolite-sex (B) associations.**

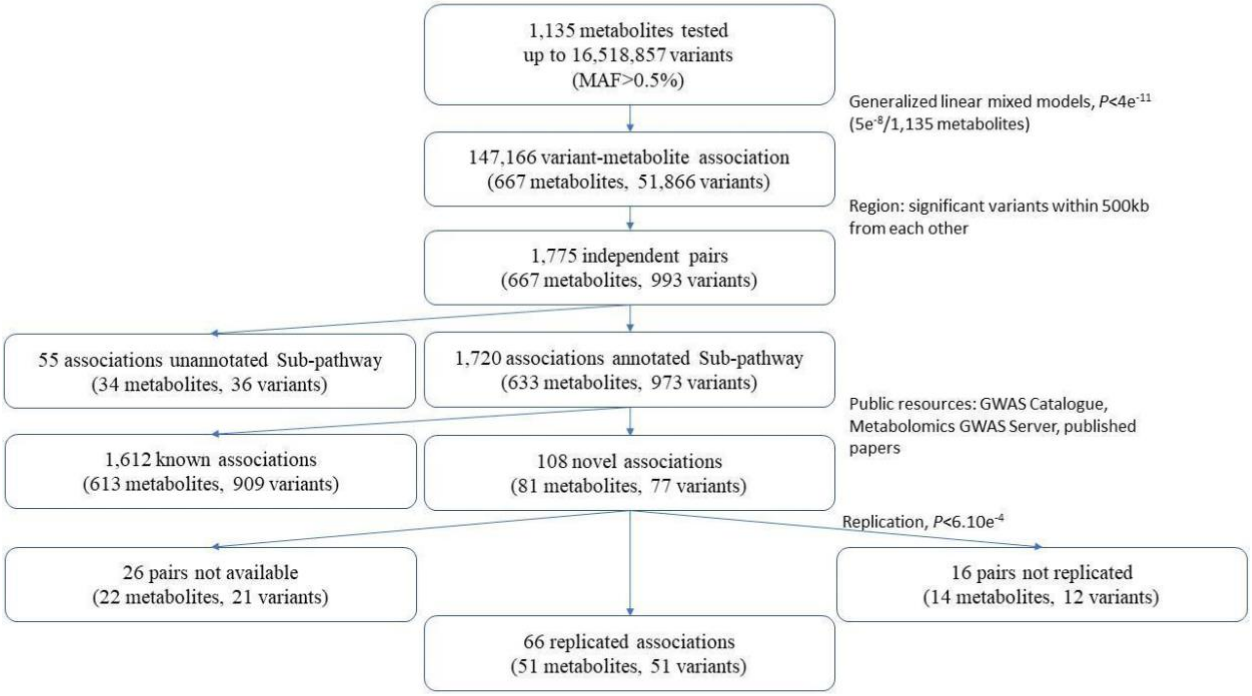


**Supplementary Figure 5. Summary of pooled metabolite QTL scanning.** We performed a single variant analysis of up to 16,518,857 variants with each of 1,135 metabolites over 16,355 participants. We set the significant association with $P \leq 4 \times 10^{-11}$. We further removed significant pairs that metabolite subpathways are not annotated. The novel association pairs were obtained by excluding known associations from public databases (see **Methods**). Only variants that were associated with a metabolite at $P \leq 6.10 \times 10^{-4}$ in our replication dataset (from Elena et al., 2013) were considered replicated.

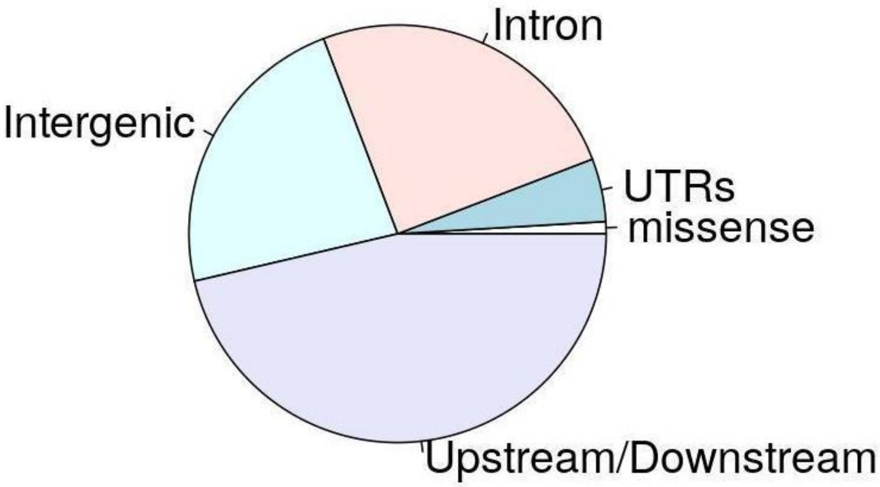


**Supplementary Figure 6.** Functional consequences of the lead variants and all significant variants within 250kb from novel variant-metabolites associations.

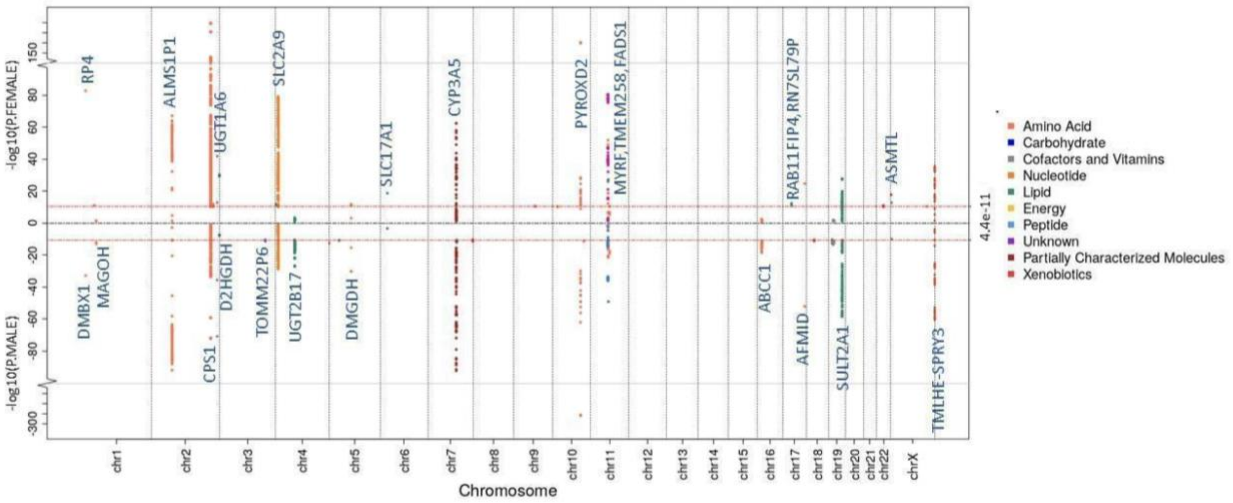

**Supplementary Figure 7. Significant associations detected from sex-stratified analyses.** Miami-Plot showing female- and male-specific association P-values from sex-stratified analyses. The upper panel shows the association from female samples and the lower panel for male samples. The x-axis represents the physical chromosome and position of each variant and the y-axis represents $-log10(P)$ from the single sex-stratified analyses. The red line indicates the genome-wide significance level on the log scale, $-log10(4.4 \times 10^{-11})$.

**A**

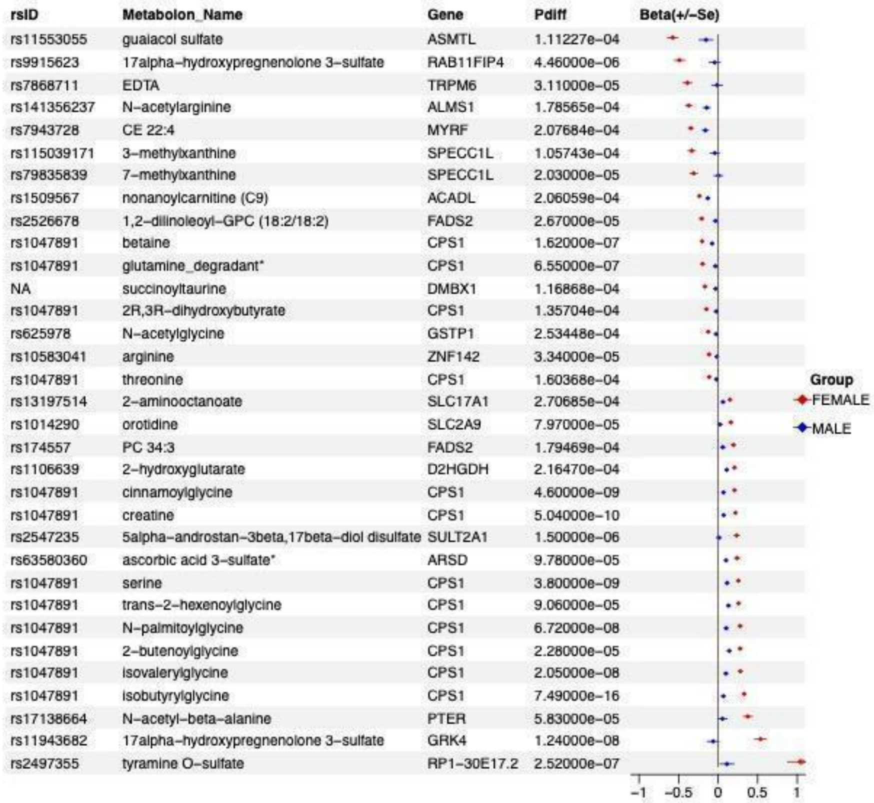
rsID Metabolon_Name Gene Pdiff Beta(+/-Se)
rs11553055 guaiacol sulfate ASMTL 1.11227e-04
rs9915623 17alpha-hydroxypregnenolone 3-sulfate RAB11FIP4 4.46000e-06
rs7868711 EDTA TRPM6 3.11000e-05
rs141356237 N-acetylarginine ALMS1 1.78565e-04
rs7943728 CE 22:4 MYRF 2.07684e-04
rs115039171 3-methylxanthine SPECC1L 1.05743e-04
rs79835839 7-methylxanthine SPECC1L 2.03000e-05
rs1509567 nonanoylcarnitine (C9) ACADL 2.06059e-04
rs2526678 1,2-dilinoleoyl-GPC (18:2/18:2) FADS2 2.67000e-05
rs1047891 betaine CPS1 1.62000e-07
rs1047891 glutamine_degradant* CPS1 6.55000e-07
NA succinoyltaurine DMBX1 1.16868e-04
rs1047891 2R,3R-dihydroxybutyrate CPS1 1.35704e-04
rs625978 N-acetylglycine GSTP1 2.53448e-04
rs10583041 arginine ZNF142 3.34000e-05
rs1047891 threonine CPS1 1.60368e-04
rs13197514 2-aminooctanoate SLC17A1 2.70685e-04
rs1014290 orotidine SLC2A9 7.97000e-05
rs174557 PC 34:3 FADS2 1.79469e-04
rs1106639 2-hydroxyglutarate D2HGDH 2.16470e-04
rs1047891 cinnamoylglycine CPS1 4.60000e-09
rs1047891 creatine CPS1 5.04000e-10
rs2547235 5alpha-androstan-3beta,17beta-diol disulfate SULT2A1 1.50000e-06
rs63580360 ascorbic acid 3-sulfate* ARSD 9.78000e-05
rs1047891 serine CPS1 3.80000e-09
rs1047891 trans-2-hexenoylglycine CPS1 9.06000e-05
rs1047891 N-palmitoylglycine CPS1 6.72000e-08
rs1047891 2-butenoylglycine CPS1 2.28000e-05
rs1047891 isovalerylglycine CPS1 2.05000e-08
rs1047891 isobutyrylglycine CPS1 7.49000e-16
rs17138664 N-acetyl-beta-alanine PTER 5.83000e-05
rs11943682 17alpha-hydroxypregnenolone 3-sulfate GRK4 1.24000e-08
rs2497355 tyramine O-sulfate RP1-30E17.2 2.52000e-07
−1 −0.5 0 0.5 1
Group
FEMALE
MALE

**B**

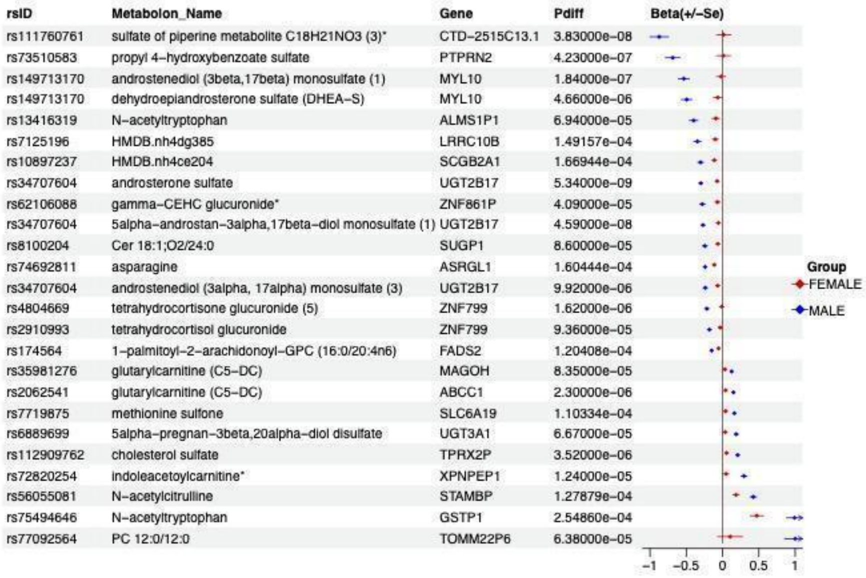
rsID Metabolon_Name Gene Pdiff Beta(+/-Se)
rs111760761 sulfate of piperine metabolite C18H21NO3 (3)* CTD-2515C13.1 3.83000e-08
rs73510583 propyl 4-hydroxybenzoate sulfate PTPRN2 4.23000e-07
rs149713170 androstenediol (3beta,17beta) monosulfate (1) MYL10 1.84000e-07
rs149713170 dehydroepiandrosterone sulfate (DHEA-S) MYL10 4.66000e-06
rs13416319 N-acetyltryptophan ALMS1P1 6.94000e-05
rs7125196 HMDB.nh4dg385 LRRC10B 1.49157e-04
rs10897237 HMDB.nh4ce204 SCGB2A1 1.66944e-04
rs34707604 androsterone sulfate UGT2B17 5.34000e-09
rs62106088 gamma-CEHC glucuronide* ZNF861P 4.09000e-05
rs34707604 5alpha-androstan-3alpha,17beta-diol monosulfate (1) UGT2B17 4.59000e-08
rs8100204 Cer 18:1;O2/24:0 SUGP1 8.60000e-05
rs74692811 asparagine ASRGL1 1.60444e-04
rs34707604 androstenediol (3alpha, 17alpha) monosulfate (3) UGT2B17 9.92000e-06
rs4804669 tetrahydrocortisone glucuronide (5) ZNF799 1.62000e-06
rs2910993 tetrahydrocortisol glucuronide ZNF799 9.36000e-05
rs174564 1-palmitoyl-2-arachidonoyl-GPC (16:0/20:4n6) FADS2 1.20408e-04
rs35981276 glutarylcarnitine (C5-DC) MAGOH 8.35000e-05
rs2062541 glutarylcarnitine (C5-DC) ABCC1 2.30000e-06
rs7719875 methionine sulfone SLC6A19 1.10334e-04
rs6889699 5alpha-pregnan-3beta,20alpha-diol disulfate UGT3A1 6.67000e-05
rs112909762 cholesterol sulfate TPRX2P 3.52000e-06
rs72820254 indoleacetoylcarnitine* XPNPEP1 1.24000e-05
rs56055081 N-acetylcitrulline STAMBP 1.27879e-04
rs75494646 N-acetyltryptophan GSTP1 2.54860e-04
rs77092564 PC 12:0/12:0 TOMM22P6 6.38000e-05
−1 −0.5 0 0.5 1
Group
FEMALE
MALE

**C**

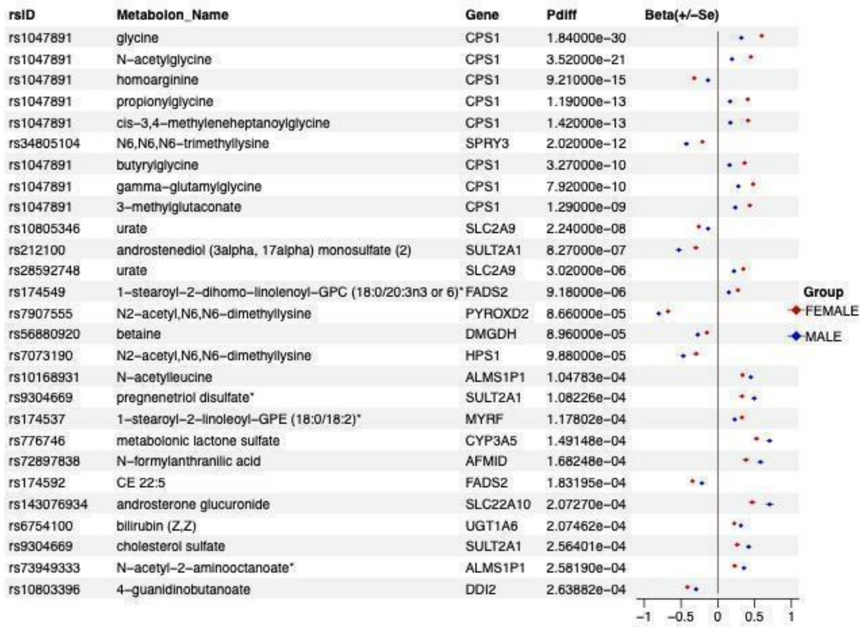


**Supplementary Figure 8. Sex-specific effect sizes of significant sex-specific SNPs on metabolite levels  from sex-stratified analyses.** The sex-stratified loci are significant in females only (**A**), males only (**B**), and both males and females but with different levels(**C**).

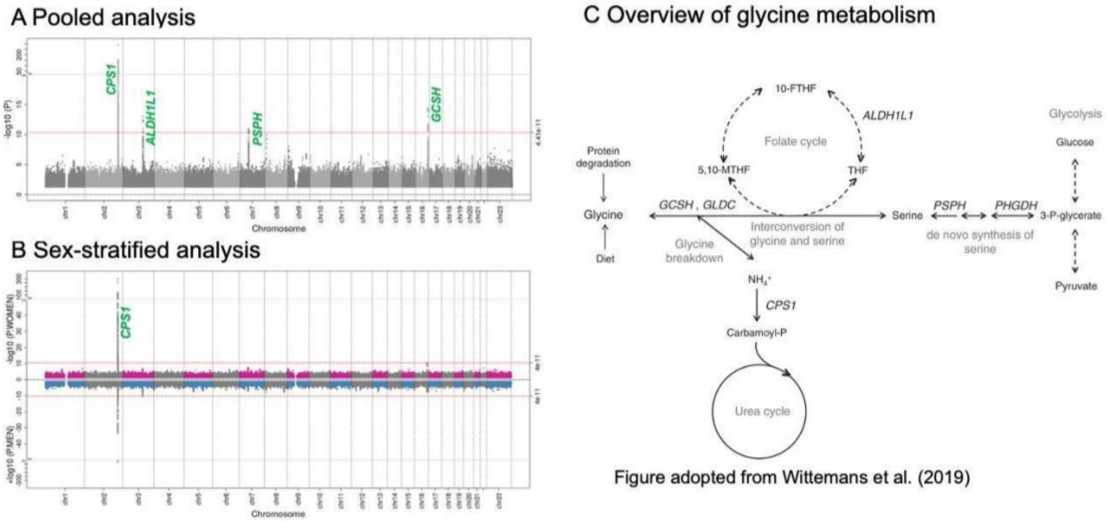


**Supplementary Figure 9. Associations of glycine metabolism from pooled metabolite QTL scanning**. A, Manhattan Plot displaying association results. The x-axis represents the physical chromosome and position of each variant and the y-axis represents $-\log_{10}(P)$ from the single variant test. The red line indicates the genome-wide significance level on the log scale, $-\log_{10}(4.4x10^{-11})$. Four genes over the significant level are highlighted. B, Miami-plot shows female- and male-specific association P-values for glycine. The upper and lower sides of the y-axis are responding to female and male separately. The red horizontal line represents the significance level of $4.4x10^{-11}$. The sex-specific genes are highlighted in green color in the miami plots.
